# RAG suppresses group 2 innate lymphoid cells

**DOI:** 10.1101/2024.04.23.590767

**Authors:** Aaron M. Ver Heul, Madison Mack, Lydia Zamidar, Masato Tamari, Ting-Lin Yang, Anna M. Trier, Do-Hyun Kim, Hannah Janzen-Meza, Steven J. Van Dyken, Chyi-Song Hsieh, Jenny M. Karo, Joseph C. Sun, Brian S. Kim

## Abstract

Antigen specificity is the central trait distinguishing adaptive from innate immune function. Assembly of antigen-specific T cell and B cell receptors occurs through V(D)J recombination mediated by the Recombinase Activating Gene endonucleases RAG1 and RAG2 (collectively called RAG). In the absence of RAG, mature T and B cells do not develop and thus RAG is critically associated with adaptive immune function. In addition to adaptive T helper 2 (Th2) cells, group 2 innate lymphoid cells (ILC2s) contribute to type 2 immune responses by producing cytokines like Interleukin-5 (IL-5) and IL-13. Although it has been reported that RAG expression modulates the function of innate natural killer (NK) cells, whether other innate immune cells such as ILC2s are affected by RAG remains unclear. We find that in RAG-deficient mice, ILC2 populations expand and produce increased IL-5 and IL-13 at steady state and contribute to increased inflammation in atopic dermatitis (AD)-like disease. Further, we show that RAG modulates ILC2 function in a cell-intrinsic manner independent of the absence or presence of adaptive T and B lymphocytes. Lastly, employing multiomic single cell analyses of RAG1 lineage-traced cells, we identify key transcriptional and epigenomic ILC2 functional programs that are suppressed by a history of RAG expression. Collectively, our data reveal a novel role for RAG in modulating innate type 2 immunity through suppression of ILC2s.

## INTRODUCTION

Atopic disorders such as atopic dermatitis (AD), asthma, and food allergy are associated with T helper type 2 (Th2) cell responses, elevated production of the type 2 cytokines interleukin(IL)-4, IL-5, and IL-13, and induction of immunoglobulin(Ig)E^1–4^.Classically, this allergic inflammatory cascade is believed to originate with antigenic stimulation of T cell receptors on adaptive T cells, which in turn results in the production of IgE from B and plasma cells capable of binding the same antigen. Indeed, presence of antigen-specific IgE reactivity is a hallmark across atopic disorders.^5^ Thus, for decades, antigen-specific adaptive Th2 cell responses have been the primary focus of investigation in the pathogenesis of atopic diseases. However, recent studies indicate that innate immune cells are sufficient to not only drive allergic pathology, but also amplify adaptive Th2 cell responses^6–9^. These studies suggest that innate immune mechanisms may play a larger role in driving atopic inflammation than previously recognized.

Innate lymphoid cells (ILCs), while lacking antigen receptors generated by recombination activating gene proteins RAG1 and RAG2 (collectively called RAG), are the innate counterparts of T cells. For example, ILC2s mirror adaptive Th2 cells in their developmental requirements, cytokine profiles, and effector functions^10^. Unlike classical T cells, ILC2s are concentrated at barrier surfaces to rapidly respond to microbial and environmental stimuli. ILC2s are key mediators of inflammatory skin conditions like AD^11–13^. Indeed, in murine models of AD-like disease, type 2 skin inflammation can still occur despite the absence of adaptive T cells, but is further reduced after depletion of ILC2s^12,13^. Furthermore, recent studies have shown that ILC2s harbor non-redundant functions in the presence of the adaptive immune system in the setting of anti-helminth immunity^14,15^. These findings suggest that ILC2 dysfunction may also uniquely contribute to the pathogenesis of atopic diseases, independent of adaptive immunity. However, the cell-intrinsic mechanisms that drive ILC2 dysregulation remain poorly understood.

ILC2s were originally discovered due to their capacity to orchestrate multiple allergic pathologies in immunocompromised mice, most notably in RAG-deficient mice that lack T and B cells^16–22^. These discoveries fundamentally redefined our understanding of allergic diseases and placed a major focus on ILC2s as potential drivers of human allergic disease. However, despite ILC2s not requiring RAG expression for their development, fate mapping studies in mice have demonstrated that up to 60% of ILC2s have historically expressed RAG1 during development^23,24^. Although previous work has described roles of RAG beyond antigen receptor recombination in developing T and B cells^25–27^ and NK cells^24^, how this developmental expression of RAG impacts ILC2s remains unclear.

By directly comparing RAG-deficient and RAG-sufficient mice, we unexpectedly found enhanced AD-like disease in RAG-deficient mice, despite the lack of adaptive lymphocytes to contribute to AD-like inflammation. Using splenocyte replenishment and bone marrow chimeras, we show that RAG suppresses ILC2 activation and expansion in a cell-intrinsic manner. Employing a RAG1-lineage reporter mouse line, we performed simultaneous single-cell multiomic RNA and ATAC sequencing to show that RAG fate-mapped ILC2s display unique transcriptional and epigenomic alterations consistent with the suppression of effector cytokine production. Collectively, our studies reveal evolutionarily conserved regulatory functions of RAG within innate lymphocytes, extending beyond the generation of antigen receptors in adaptive lymphocytes.

## RESULTS

### RAG deficiency leads to expansion and activation of ILC2s

AD-like disease can be elicited in the skin of mice with repeated application of the topical vitamin D analog calcipotriol (MC903)^28^. Although it has been previously demonstrated that MC903 can induce AD-like disease in RAG-deficient mice that lack T and B cells, in part via ILC2 activation^12,13^, the relative contributions of ILC2s and the adaptive lymphocyte compartment have not been rigorously evaluated. We hypothesized that the presence of Th2 cells, in addition to ILC2s, would lead to enhanced AD-like disease in an additive fashion. In testing this, we evaluated both RAG1-sufficient wild-type (WT) mice and RAG1-deficient *Rag1^-/-^* mice in the setting of AD-like disease (**Fig. 1A**). Unexpectedly, we observed that *Rag1^-/-^* mice developed increased ear skin thickness (**Fig. 1B**) and increased absolute numbers and proportion of ILC2s in the skin-draining lymph nodes (sdLNs) compared to control WT mice (**Fig. 1C,D; S1A,B**). Furthermore, a larger proportion of ILC2s from *Rag1^-/-^* mice exhibited production of both IL-5 (**Fig. 1E, S1C**) and IL-13 (**Fig. 1F, S1C**). Our findings indicated that RAG1 deficiency results in paradoxically worse AD-like disease in association with enhanced ILC2 expansion and activation.

**Figure 1.**
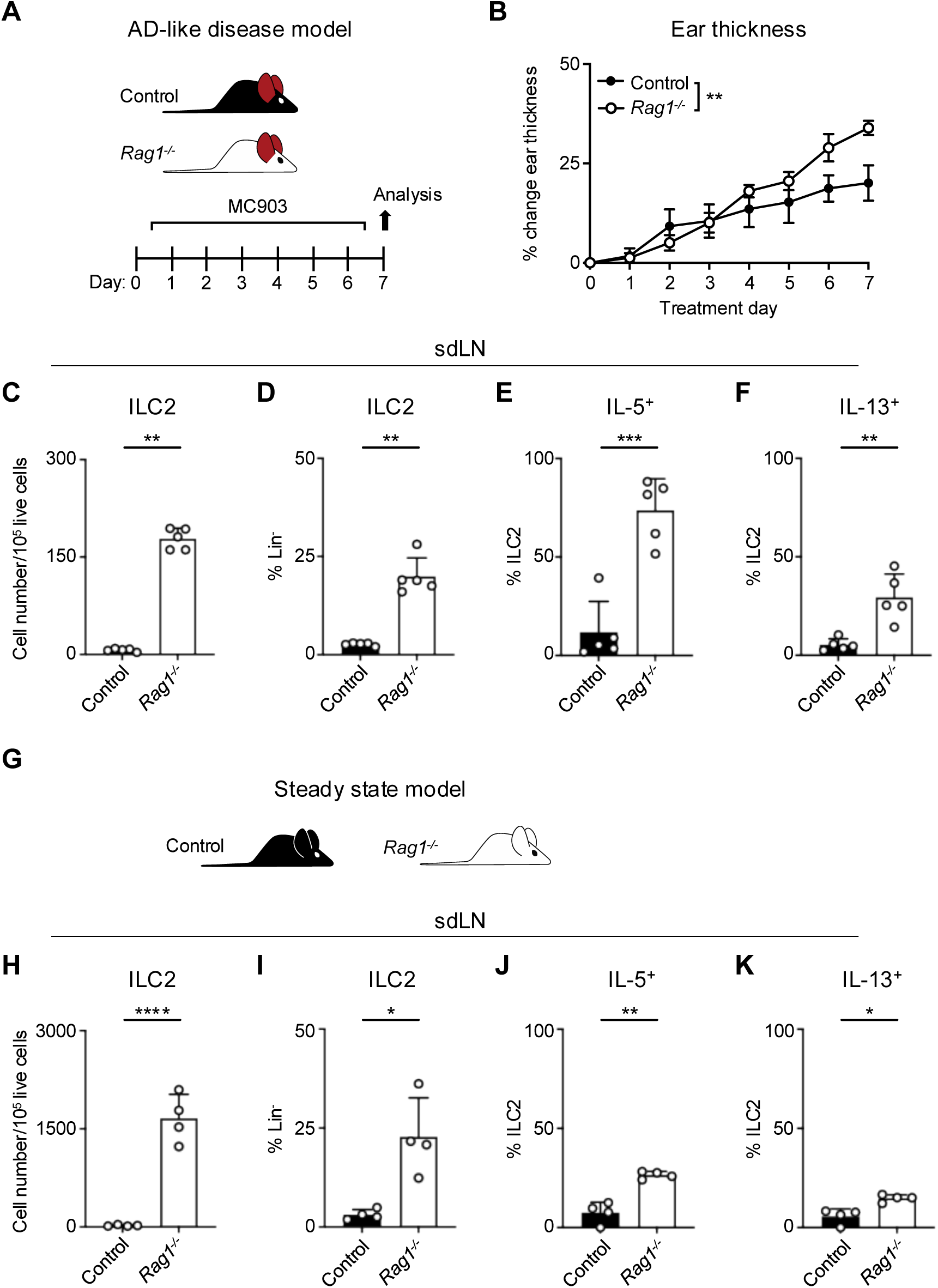
RAG deficiency leads to expansion and activation of ILC2s during inflammation and at steady state. **A)** Experimental schematic of AD-like disease. WT (Control) B6 mice or *Rag1^-/-^* mice treated topically to the inner surface of each ear with 2 nmol MC903 in 10 μL ethanol vehicle daily for 7 days develop AD-like inflammation. **B)** Ear thickness measured daily in AD-like inflammation. Data representative of at least 2 independent experiments, 5 mice/group. ** P < 0.01 by 2-way ANOVA with Sidak’s multiple comparisons test, day 7. **C)** Total number ILC2s normalized to 10^5^ live cells and **D**) ILC2 proportion of CD90^+^, Lin^-^ cells (Lin^-^ defined as CD3^-^, CD5^-^, CD11b^-^, CD11c^-^, CD19^-^, NK1.1^-^, and FcεR1^-^) determined to be ILC2s (IL-33R^+^) in skin-draining lymph nodes (sdLN) from WT or *Rag1^-/-^* mice with AD-like ear inflammation. Percent ILC2 from sdLN in AD-like disease following PMA/iono stimulation positive for **E)** IL-5 or **F)** IL-13 staining. **G)** Schematic of steady state analysis of sdLN from WT (Control) or *Rag1^-/-^* mice. **H)** Total number ILC2s normalized to 10^5^ live cells and **I**) ILC2 proportion of steady state sdLN CD90^+^, Lin^-^ cells determined to be ILC2s as in **(C,D)**. Percent ILC2 from sdLN in steady state following PMA/iono stimulation positive for **J)** IL-5 or **K)** IL-13 staining. **C-F; H-K)** Data representative of at least 2 independent experiments, 4-5 mice/group * P < 0.05, ** P < 0.01, *** P < 0.001 by two-tailed Welch’s t test. All data represented as mean with standard deviation.

To determine whether this phenomenon was specific to AD-like pathological conditions, we next examined the sdLNs in *Rag1^-/-^* and lymphocyte-sufficient *Rag^+/-^* littermate control mice in the absence of disease (**Fig. 1G**). We found that the absolute number and frequency of ILC2s was increased at steady state in *Rag1^-/-^* sdLNs (**Fig. 1H,I**) and that a higher proportion of these ILC2s produced both IL-5 (**Fig. 1J**) and IL-13 (**Fig. 1K**) compared to WT controls following ex vivo stimulation and intracellular cytokine staining. The RAG recombinase requires both RAG1 and RAG2 components to successfully rearrange a functional antigen receptor in adaptive lymphocytes^29^. Thus, to test whether our findings are specific to RAG1, or related to function of the overall RAG complex, we similarly examined the steady-state profile of ILC2s in *Rag2^-/-^* mice (**Fig. S2A**). Deficiency of RAG2 led to an expansion of ILC2s in the sdLNs (**Fig. S2B,C**) and increased proportions of ILC2s expressing IL-5 (**Fig. S2D**) and IL-13 (**Fig. S2E**) similar to *Rag1^-/-^*mice. Collectively, these findings suggest that the RAG recombinase modulates ILC2 function at steady state and during type 2 inflammation. However, whether the hyperactive ILC2 phenotype is due to a cell-intrinsic process or simply due to the absence of T and B cells was unclear.

### ILC2 suppression by RAG is cell intrinsic

Given the importance of the adaptive lymphocyte compartment in shaping the secondary lymphoid organ repertoire, we next wanted to examine whether the presence of adaptive lymphocytes could restore ILC2 homeostasis in RAG-deficient mice. To test this, we created splenocyte chimera mice by reconstituting both *Rag1^-/-^*and control WT mice with splenocytes containing T and B cells from WT donor mice (**Fig. 2A**). We first assessed the overall level of immune reconstitution in the recipient mice and found fully restored proportions of CD4^+^ (**Fig. S3A**) and CD8^+^ (**Fig. S3B**) T cells in the spleens of recipient *Rag1^-/-^* mice, although B cell numbers remained significantly lower than in WT mice (**Fig. S3C**). Upon induction of AD-like disease, we found that the *Rag1^-/-^*mice still exhibited increased ear skin thickness (**Fig. 2B**), enhanced expansion of ILC2s (**Fig. 2C,D**), and increased proportions of ILC2s expressing IL-5 (**Fig. 2E**) and IL-13 (**Fig. 2F**) in the sdLNs. Interestingly, we found significantly higher proportions of eosinophils in the spleens of *Rag1^-/-^* recipient mice (**Fig. S3D**), possibly reflecting the increased IL-5 production we observed from ILC2s. These findings indicate that the mere introduction of exogenous T and B cells is not sufficient to suppress ILC2 dysregulation in the setting of RAG deficiency.

**Figure 2.**
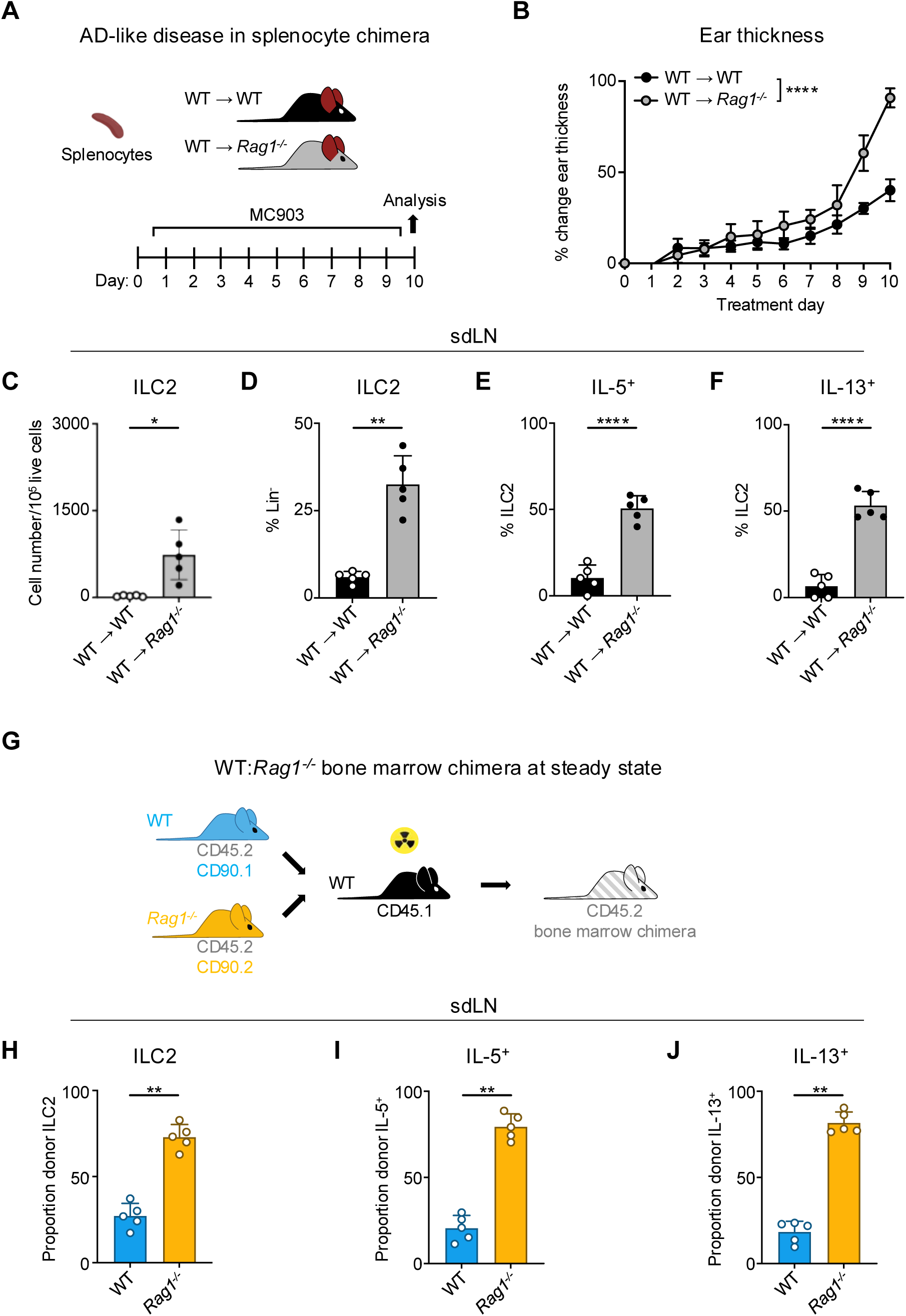
Homeostatic expansion and activation of RAG-deficient ILC2s is cell intrinsic. **A)** Experimental schematic of AD-like disease in splenocyte chimera experiment. WT B6 or *Rag1^-/-^*mice received WT splenocytes and developed AD-like inflammation after subsequent topical treatment with 2 nmol MC903 in 10 μL ethanol vehicle to each ear daily for 10 days. **B)** Ear thickness measured daily in AD-like inflammation. Data representative of 2 independent experiments, 4-5 mice per group. **** P <0.0001 by 2-way ANOVA with Sidak’s multiple comparisons test, day 10. **C)** Total number ILC2s normalized to 10^5^ live cells and **D**) ILC2 proportion of CD90^+^, Lin^-^ cells (Lin^-^ defined as CD3^-^, CD5^-^, CD11b^-^, CD11c^-^, CD19^-^, NK1.1^-^, and FcεR1^-^) determined to be ILC2s (IL-33R^+^). Percent ILC2 from sdLN in splenocyte chimera mice with AD-like disease after PMA/iono stimulation positive for **E)** IL-5 or **F)** IL-13 staining. **G)** Schematic of bone marrow chimera experiment. Equal quantities of bone marrow cells from *Rag1^-/-^* (CD45.2, CD90.2 - orange) and WT (CD45.2, CD90.1 - blue) C57Bl/6J donor mice were used to reconstitute the immune systems of irradiated recipient WT (CD45.1 - black) C57Bl/6J mice. **H)** Proportion of donor (CD45.2^+^) ILC2 defined as in (C) in sdLN by donor source (CD90.1^+^ - WT, CD90.2^-^ - *Rag1*^-/-^). Proportion of Lin^-^ ILCs by donor source positive for **I)** IL-5 and **J)** IL-13 following PMA/iono stimulation and cytokine staining. **C-F)** Data representative of at least 2 independent experiments, 4-5 mice per group. ** P < 0.01, **** P < 0.0001 by two-tailed Welch’s t test. **H-J)** Data representative of at least 2 independent experiments, 4-5 mice per group. ** P < 0.01 by two-tailed ratio means paired t test. All data represented as mean with standard deviation.

To further test whether this phenotype is mediated by cell-intrinsic RAG expression, we next generated mixed bone marrow (BM) chimeras. We harvested BM from congenic CD90.1^+^ WT and CD90.2^+^ *Rag1^-/-^* donor mice on a CD45.2^+^ background and reconstituted sub-lethally irradiated CD45.1^+^ congenic WT recipients with a 50:50 mixture of WT:*Rag1^-/-^* BM (**Fig. 2G**). After confirming reconstitution of donor immune cells in the sdLN (**Fig. S4A,D**), we examined the frequency and activity of ILC2s in the sdLNs based on whether they originated from WT (CD90.1^+^) or *Rag1^-/-^*(CD90.2^+^) donors (**Fig. S4B,C**). Strikingly, of the total donor ILC2s, the majority were derived from *Rag1^-/-^* donors (**Fig. 2F,G**). This was not due to differences in overall donor reconstitution, since measuring all Lin^-^ cells revealed WT donor cells outnumbered those from *Rag1^-/-^* donors (**Fig. S4E**). Of the total IL-5- (**Fig. 2H,I,K**) and IL-13-expressing (**Fig. 2 H,J,L**) ILCs, the majority were also derived from *Rag1^-/-^* donors. Taken together, these data suggest that cell-intrinsic RAG activity in ILC2s may limit their capacity to expand and become activated.

### A history of RAG expression marks a subpopulation of ILC2s in the skin draining lymph node

In contrast to resting naïve T cells, ILC2s resemble activated Th2 cells at steady state based on their transcriptomic and epigenomic profiles^30,31^. While both T cells and ILC2s exhibit historical RAG expression^23^, they do not actively express the protein in their mature state^32^. Taken together, these findings provoke the hypothesis that ILC2s are regulated by RAG early in development to imprint alterations that influence their activity as mature cells. To distinguish ILC2s as either having a history of RAG expression or not, we utilized a RAG lineage tracing system, whereby a *Rag1^Cre^*mouse was crossed to a reporter mouse expressing tandem dimer red fluorescent protein (tdRFP) in a Cre-dependent manner from the *Rosa26* locus (**Fig. 3A**)^24,33^. This system allowed us to compare RAG-experienced (RAG^exp^) and RAG-naïve (RAG^naïve^) lymphoid cells, including ILC2s, simultaneously originating from the same immunocompetent host, thus removing confounders inherent in knockout and chimera experiments. Analysis of sdLN from the reporter mice revealed that nearly all CD4^+^ T cells, CD8^+^ T cells, and B220^+^ B cells expressed tdRFP (positive history of Cre expression from the *Rag1* locus), consistent with the known requirement of RAG expression for their development (**Fig. 3B-D,G**). We also examined NK cells, since certain subsets of NK cells are known to express RAG during their development^24^, and we observed that 60% of NK cells were tdRFP^+^, similar to previous findings (**Fig. 3E,G**)^24^. In the ILC2 population of the sdLN, around 50% were tdRFP^+^ (**Fig. 3F,G**), similar to proportions of RAG fate-mapped ILC2s previously observed in the fat^24^ and lung^23,24^. These findings demonstrate that there are heterogeneous populations of ILC2s marked by differential tdRFP^+^ fate mapping. Importantly, this provided us with the possibility to profile these different ILC2 subsets based on *Rag1^Cre^*-activated expression of tdRFP.

**Figure 3.**
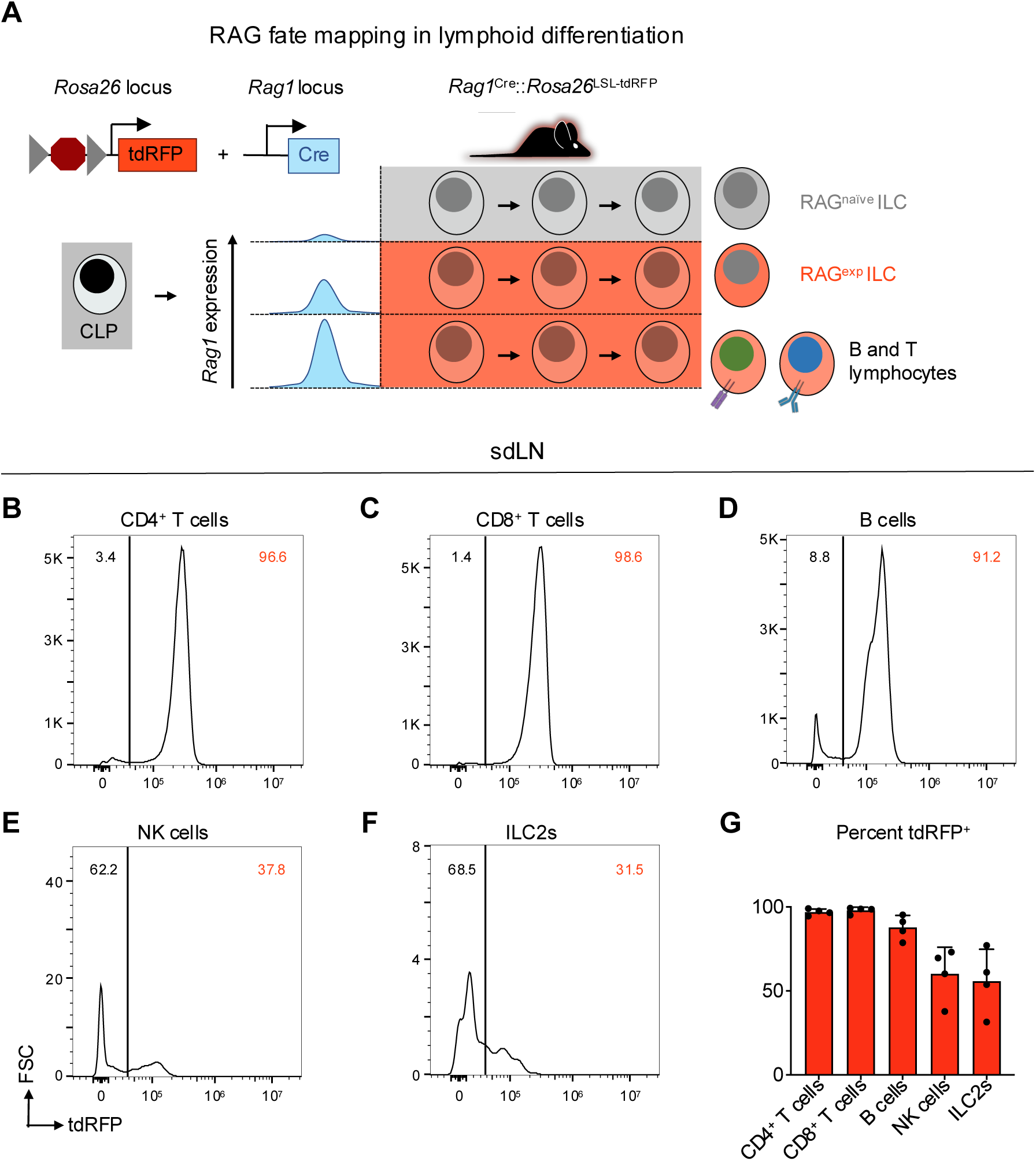
A history of RAG expression marks a population of ILC2s in the sdLN. **A)** Schematic of RAG fate mapping in the lymphoid cell compartment using reporter mice expressing Cre-inducible tandem dimer red fluorescent protein (tdRFP) from the *Rosa26* locus crossed to mice expressing Cre recombinase from the *Rag1* locus. **B-F)** Histograms of tdRFP signal in CD45^+^ sdLN cells by cell type for **B)** CD4^+^ T cells (B220^-^, CD3^+^, CD4^+^), **C)** CD8^+^ T cells (B220^-^, CD3^+^, CD8^+^), **D)** B cells (MHCII^+^, B220^+^), **E)** NK cells (B220^-^, CD3^-^, CD4^-^, CD8^-^, CD49b^+^, NK1.1^+^), **F)** ILC2s (B220^-^, CD3^-^, CD4^-^, CD8^-^, CD49b^-^, NK1.1^-^, CD11b^-^, CD11c^-^, SiglecF^-^, CD90^+^, KLRG1^+^ or ICOS^+^ or IL-33R^+^), **F)** quantification of tdRFP^+^ proportion of each cell type.

### Multiomic profiling enhances the detection of rare tissue-specific ILC2s

Transient RAG expression early in lymphoid development leads to well-characterized, durable effects on B and T cell development and function mainly through successful genomic rearrangement of antigen receptors. Yet, our data indicate that RAG expression also imprints phenotypic changes on ILC2s, which can develop independently of functional antigen receptors, provoking the hypothesis that RAG expression may affect broader epigenomic and transcriptional programs. Furthermore, our data indicate that the impact of RAG on ILC2 function has implications for AD-like skin inflammation, suggesting a persistent effect that modulates phenotypes of type 2 inflammation.

To test these hypotheses, we performed combined single-nuclei RNA sequencing (snRNA-seq) and ATAC sequencing (snATAC-seq) of sdLN cells from RAG fate-mapped mice at steady state and in the setting of AD-like disease (**Fig. S5A,B**). Because fluorescence-activated cell sorting (FACS) can cause physical stress, cell loss, and contamination, which can introduce unwanted perturbations in target cells, instead we utilized gentle initial negative selection by magnetic activated cell sorting (MACS) to remove most B and T cells and monocytes prior to sequencing (**Fig. S5A**). This allowed us to enrich for innate immune cell populations prior to sequencing. Further, we used the gene encoding tdRFP as a barcode to differentiate between RAG^exp^ (RAG fate map-positive) and RAG^naïve^ (RAG fate map-negative) ILC2s at the single-cell level (**Fig. 5A**). The multiomic data was analyzed using recently developed pipelines in Cell Ranger, Seurat^34–36^, and Signac^37^, and sequenced cells were further filtered computationally to enrich for ILCs, as in previous studies (see methods)^38^.

In addition to gene expression (GEX) information derived from snRNA-seq (**Fig. 4A**), we calculated a “gene activity” (GA) score based on chromatin accessibility at gene loci (**Fig. 4B**) from the corresponding snATAC-seq dataset^37^. Clustering the cells with each data subset alone and in combination using weighted nearest neighbor (WNN) analysis, we identified six clusters **(Fig. 4C)** that demonstrated consistent differences in cellular markers based on both metrics of GEX and GA (**Fig. 4D,E; Table S1**). Additionally, top markers for each cluster clearly differentiated each cell type (**Fig. 4F**). Despite successful ILC2 enrichment via MACS depletion for lineage markers and computational filtering (see methods), our data set included non-ILC2 populations determined by gene expression to be T cells, dendritic cells, B cells, and NK cells (**Fig. 4A,F; Table S1**), which allowed for broader multidimensional comparisons while studying this ILC2-enriched data set.

**Figure 4.**
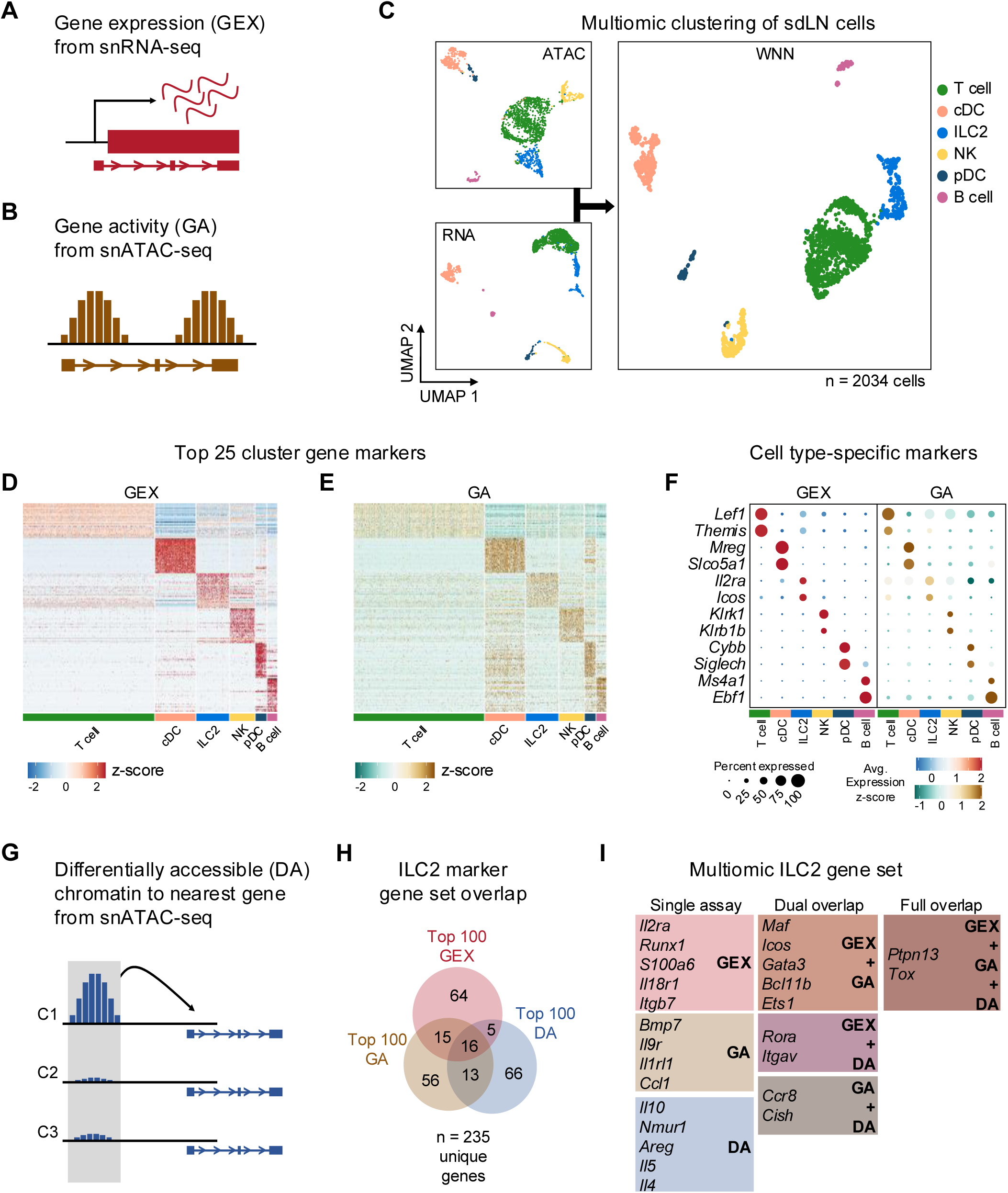
Multiomic analysis of ILC2s through single nuclei sequencing of the sdLN. **A)** Schematic of the gene expression (GEX) assay derived from snRNA-seq data. **B)** Schematic of the gene activity (GA) assay, representing estimated transcription scores derived from snATAC-seq data using Signac. **C)** UMAP visualizations of independent analyses of RNA-seq and ATAC-seq data for 2034 sdLN cells after dimensional reduction and clustering combined using weighted nearest neighbor (WNN) analysis in Seurat. Cluster identities are color coded consistently throughout the following panels. Heatmaps of **D)** top 25 GEX marker genes and **E)** top 25 GA marker genes identified for each cluster. See **Table S1** for full lists of genes. **F)** Dotplots comparing selected marker genes for each cluster between the GEX and GA assays, with emphasis on known cell type-specific markers. **G)** Schematic of differentially accessible (DA) chromatin assay, which finds the nearest gene to any peak calculated to be differentially open in a particular cell cluster. See **Table S2** for full lists of top 25 DA cluster markers. **H)** Overlap of top 100 markers for the ILC2 cluster from the GEX, GA, and DA assays. See **Table S3** for top 100 DA peaks and distances to nearest genes and **Table S4** for full list of top 100 ILC2 markers. **I)** Selected genes from the ILC2 gene set for each assay individually and for overlaps.

To further complement the GEX and GA assays, we utilized another method of detecting cell-specific marker genes, whereby chromatin regions that are differentially accessible (DA), or open, in each cluster could be linked by their physical proximity to specific genes (**Fig. 4G, S6**; **Table S2**; see methods). Comparing the top 100 GEX, GA, and DA markers in the ILC2 population, we identified a multiomic ILC2 signature of 235 unique genes (**Fig. 4H; Table S3,4**). While there was some overlap between each respective set, the multiomic approach enabled more extensive identification of an ILC2 gene program than through either snRNA-seq or snATAC-seq alone. The analysis revealed a variety of canonical ILC2-associated genes specific to the ILC2 cluster (**Fig. 4I, S6,7**) including the ILC2-activating cell surface receptors *Icos*^39,40^, *Il2ra*^41,42^, *Il18r1*^38,43,44^, *Nmur*^14,15,45–47^, and *Il1rl1* (encoding the receptor for IL-33)^11,12,48,49^, as well as transcription factors such as *Gata3*^19,50–53^, *Bcl11b*^54–57^, *Maf*^19,58,59^, *Ets1*^60^, and *Rora*^38,51,61–63^, all previously shown to be important for ILC2 development and/or function.

Expression of some secreted proteins can be difficult to capture in droplet-based snRNA-seq experiments due to low transcript levels and relatively shallow sequencing depth. With the addition of the complementary GA and DA assays from snATAC-seq, our analysis identified *Il5* (**Fig. 4I, S6B**), a canonical ILC2 cytokine, in the DA assay, while in the GA assay we found *Bmp7*^64,65^, which has been shown to be secreted by ILC2s to influence browning of adipose tissue. Additionally, we identified the secreted chemokine *Ccl1* as an ILC2 marker gene^66–69^, which along with its cognate receptor *Ccr8* (also an ILC2 marker in our analysis)^44,67,70^, participates in a feed-forward circuit to drive ILC2 recruitment and expansion^71^. Thus, our findings demonstrate how genetic barcoding, combining transcriptomic and epigenomic analyses, and cross-validation across many published studies can yield new insights while providing internal control measures to elevate the rigor, robustness, and confidence of identifying gene signatures of rare populations such as ILC2s at the single-cell level.

### ILC2s with a history of RAG expression are epigenomically suppressed

As noted above, barcoding the ILC2s afforded the opportunity to transcriptionally and epigenomically profile ILC2s under identical developmental conditions by dividing the ILC2 cluster into RAG^exp^ and RAG^naïve^ populations. We hypothesized that RAG^exp^ ILC2s would have a distinct transcriptional profile compared to ILC2s without any history of RAG expression. To test this, we calculated differentially expressed genes (DEGs) for the ILC2 cluster by *Rag1* fate-map status. Genes with higher expression in RAG^exp^ cells relative to RAG^naïve^ cells had positive fold change values, and vice versa, with genes relatively increased in RAG^naïve^ cells having negative values (**Fig. S8A, Table S5**). Using gene set enrichment analysis (GSEA)^72,73^ on the ranked list of DEGs, we found that gene sets generally representing lymphocyte activation and differentiation were suppressed in RAG^exp^ ILC2s compared to RAG^naïve^ ILC2s (**Fig. S8B,C, Table S6**), consistent with our previous observations that ILC2s are expanded and more activated in RAG-deficient mice relative to WT mice.

We next employed newly described methodologies^37,74^ that quantify associations between open chromatin peaks and the expression of nearby genes to describe the functional regulomes of both RAG^exp^ and RAG^naïve^ ILC2s (**Fig. 5B**). In this analysis, each ATAC peak can be linked to multiple genes, and each gene to multiple peaks, generating a list of “gene-to-peak links” or GPLs (see methods). For each gene, we interpreted the number of corresponding GPLs as a quantitative representation of the regulome activity for that gene. Considering RAG^exp^ and RAG^naïve^ cells as two separate populations, we generated two lists of GPLs (**Table S7**) defining functional regulomes for each population. We focused our analysis on the functional regulomes of ILC2s by filtering the GPL lists for the 235 unique ILC2 genes identified in our multiomic gene set (**Fig. 4H; Table S8**), then calculated the difference in GPLs between RAG^exp^ and RAG^naïve^ cells for each gene and sorted them; genes displaying greater numbers of GPLs in the RAG^naïve^ population are at the top, and genes with more GPLs in the RAG^exp^ population are at the bottom (**Fig. 5C; Table S9**).

**Figure 5.**
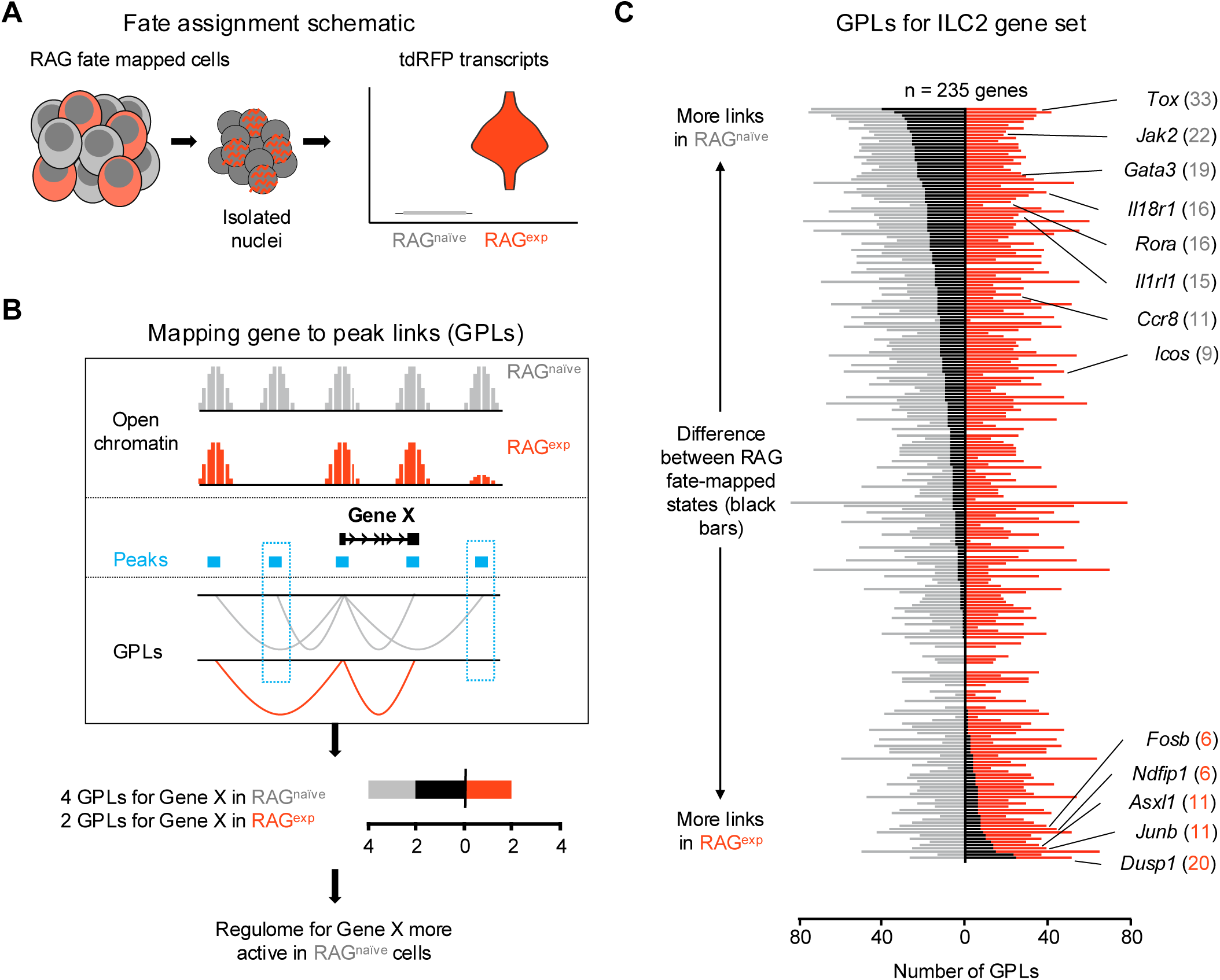
A history of RAG expression imprints transcriptomic and epigenomic modulation of ILC2 gene programs. **A)** Schematic of transcriptional RAG fate mapping. Sequenced cells from the RAG fate map mouse (see Figure 3A) transcribe tdRFP only after Cre is expressed from the *Rag1* locus. Cells were assigned as either having a history of RAG expression (RAG^exp^ - tomato red) or not (RAG^naïve^ - dark gray) based on detection of tdRFP transcript in the RNA-seq data. **B)** Schematic of mapping gene to peak links (GPLs). The LinkPeaks function of Signac (see methods) calculates significant correlations between open chromatin at defined peaks (teal bars) and nearby gene expression. These links represent inferred epigenomic-transcriptomic regulation, or "regulomes" based on the correlated snRNA- and snATAC-sequencing data. After calculating GPLs separately for each population (gray for RAG^naïve^ and tomato red for RAG^exp^), GPLs found in only one group, but not the other, can be identified (teal boxes). The difference in GPLs based on RAG experience for any given gene (e.g. Gene X) can be visualized on a bar graph, with the number of GPLs for RAG^naïve^ (gray - left) and RAG^exp^ (red - right) plotted, with the difference overlaid as a black bar. **C)** GPLs calculated as in (B) for the multiomic ILC2 gene set identified in Figure 4H and **Table S4**). All identified GPLs are listed in **Table S7**, while ILC2 GPLs are listed in **Table S8**. Genes are sorted from more links identified in the RAG^naïve^ population at top to more links identified in the RAG^exp^ population at bottom. Select genes are labeled. Full ranked list by difference in GPLs is available in **Table S9**.

We found that the ILC2 marker genes segregating toward the top of this list, corresponding to enhanced epigenomic activity in the RAG^naïve^ cells, tended to be genes previously identified to play positive roles in development, expansion, and activation of lymphoid cells. These included transcriptional regulators such as *Tox*^75–78^, *Rora*^38,51,61–63^, *Maf*^19,58,59^, and *Gata3*^19,50–52^, which are involved in early differentiation of both ILCs and lymphocytes (**Fig. S9A**). Indeed, epigenetic activation of the *Gata3* locus is recognized to play a critical role in development of both ILC2s^79^ and Th2 cells^80,81^. Additionally, surface receptors known to drive ILC2 activation upon stimulation including *Il18r1*^38,43,44^, *Il1rl1*^21,38,43,82,83^, and *Icos*^39,40^ had increased functional regulome activity in RAG^naïve^ ILC2s. In contrast, genes with more GPLs in the RAG^exp^ ILC2s tended to be associated with suppressive functions. For example, *Ndfip1* (**Fig. S9B**) encodes a regulatory protein that enhances activity of the ubiquitin ligase ITCH to negatively regulate inflammation^84,85^ and has been associated with asthma risk in GWAS studies^86^. *Dusp1* partially mediates glucocorticoid effects through its ability to negatively regulate inflammation^87,88^, is associated with eczema by GWAS^89^, and has recently been shown to mark an anti-inflammatory set of ILCs^69^. Last, *Asxl1* encodes a tumor suppressor that inhibits clonal hematopoiesis through its epigenomic regulatory effects in both mice and humans^90–93^. Collectively, our GPL analysis stratifies the ILC2 gene signature based on RAG experience, where genes associated with ILC2 expansion and activation are poised in RAG^naïve^ cells, while genes associated with suppressive effects are poised in RAG^exp^ cells.

We expanded our multiomic analysis to infer information about transcription factor (TF) activity from open chromatin regions in our snATAC data. We used the chromVAR package^94^, which finds known TF binding motifs in open chromatin regions in each cell, to identify TF motifs enriched in each cell cluster (**Fig. S10A,B**, **Table S10**). The enriched motifs were consistent with known functional roles of associated TFs in each cell type. For example, in the NK cell cluster we found enriched motifs recognized by the TFs EOMES and T-bet (encoded by *Eomes* and *Tbx21*, respectively), which are critical for development of NK cells^95^. A limitation of this analysis is that while TF motif accessibility can be inferred from open chromatin in snATAC data, which TFs are bound to the identified accessible sites is not known. We reasoned that complementary gene expression information from our multiomic data could mitigate this limitation in part by comparing accessibility of TF binding motifs to expression levels of corresponding TFs (**Fig. S10C)**. Indeed, motifs for both RORα and RORγ (encoded by *Rora* and *Rorg*, respectively), which share a common DNA binding 5’-AGGTCA-3’ half site, have similar calculated accessibilities in both the ILC2 and NK cell clusters. Yet only *Rora* is expressed at appreciable levels, and only in ILC2s, consistent with its critical role in ILC2 development^61,62^. In contrast, ILC2 development is not dependent on *Rorg* expression, and neither RORα nor RORγ plays a major role in NK cells. Taken together, this analysis confirms the known role of *Rora* in ILC2s and highlights how matched multiomic chromatin accessibility and gene expression data can clarify ambiguities inherent in TF enrichment analyses.

The broad effects of RAG expression on ILC2 transcriptional regulomes we observed (**Fig. 5C**) led us to hypothesize that distinct cohorts of TFs may contribute to the differences observed between RAG^naïve^ and RAG^exp^ ILC2s. To test this hypothesis, we analyzed the open chromatin regions in GPLs unique to each RAG fate mapped ILC2 population using the FindMotifs function in Signac^37^, which returns a ranked list of enriched motifs corrected for background presence of each motif in all cells. In both RAG^naïve^ and RAG^exp^ ILC2s, we identified enriched TF motifs (**Fig. S10D, Table S11**) that are GC rich regions recognized by a large family of C2H2 zinc finger TFs, particularly the Krüppel-like factors (KLFs), which are well-established as key regulators of lymphocyte development^96,97^. Given the strong sequence similarities of the identified TF motifs, we turned to the matched gene expression data to clarify which TFs may be available to engage the accessible binding sites. Of the eleven unique TFs identified, only six were detected in the gene expression assay (**Fig. S10E**). We observed much higher expression of *Klf2*, *Klf6*, and *Klf12* in Rag^exp^ ILC2s compared to RAG^naïve^ ILC2s (**Fig. S10E**). Notably, all three of these TFs have been associated with reduced cellular proliferation and/or activation^98–100^. *Klf2* expression plays a key role in T cell quiescence^101,102^, and both *Klf2* and *Klf6* were recently identified as markers of “quiescent-like” skin resident ILCs^69^. In contrast, although detected in a smaller fraction of cells in our data, we found *Klf7* expression was higher in RAG^naïve^ ILC2s compared to Rag^exp^ ILC2s (**Fig. S10E**). Increased expression of *Klf7* has been shown to enhance survival of early thymocytes and is a predictor of poor outcome in acute lymphoblastic leukemia^103^. Collectively, these findings link the relatively activated or suppressed epigenomic and transcriptomic states of RAG^naïve^ and Rag^exp^ ILC2s, respectively, to distinct cohorts of homeostatic TFs.

### A history of RAG expression modulates ILC2 epigenomes at steady state and in AD-like inflammation

Although our GPL and TF analyses revealed a suppressive effect of RAG expression on ILC2 gene programs, we did not account for the additional variable of disease state in the initial analysis. To test whether RAG expression promotes a suppressive epigenomic program in ILC2s that is durable in the setting of inflammation, we first recalculated GPLs after splitting our dataset by both history of RAG expression (naïve vs. experienced) and disease (steady state vs. AD-like disease) to yield four lists of GPLs (**Fig 6A; Table S12**). When we examined the intersection, or overlap, of peaks from ILC2 GPLs (**Table S13**), several notable patterns emerged (**Fig. 6B**). First, the largest set of peaks was shared by all RAG^naïve^ cells (gray bar), regardless of disease state, with the next largest peak sets belonging to either steady state or AD-like disease in the RAG^naïve^ cells. Second, there was a large set of peaks shared by all RAG^exp^ cells (red bar). Third, the intersections corresponding to disease states (steady state – yellow, AD-like disease – dark red), had relatively few unique peaks. These findings suggest that early exposure to RAG expression plays a larger role in modulating the epigenomic signature of the ILC2 gene program than exposure to disease. To confirm that the patterns we observed represent a specific effect of RAG expression on the ILC2 gene program, we performed the same analysis on GPL peaks for all genes in the dataset (**Fig. S11**). In contrast to the ILC2 gene set, the majority of GPL peaks for all genes was shared among all cell populations, consistent with epigenomic regulation of most genes being minimally affected by either RAG expression or AD-like disease. Last, in the ILC2 gene set analysis, we noted a set of poised peaks shared by all RAG^naive^ cells and RAG^exp^ cells in the setting of AD-like disease, but not with RAG^exp^ cells at steady state (blue bar, **Fig. 6B**). We reasoned that this condition might capture some genomic loci that are suppressed by a history of RAG expression at steady state but are induced during inflammation.

**Figure 6.**
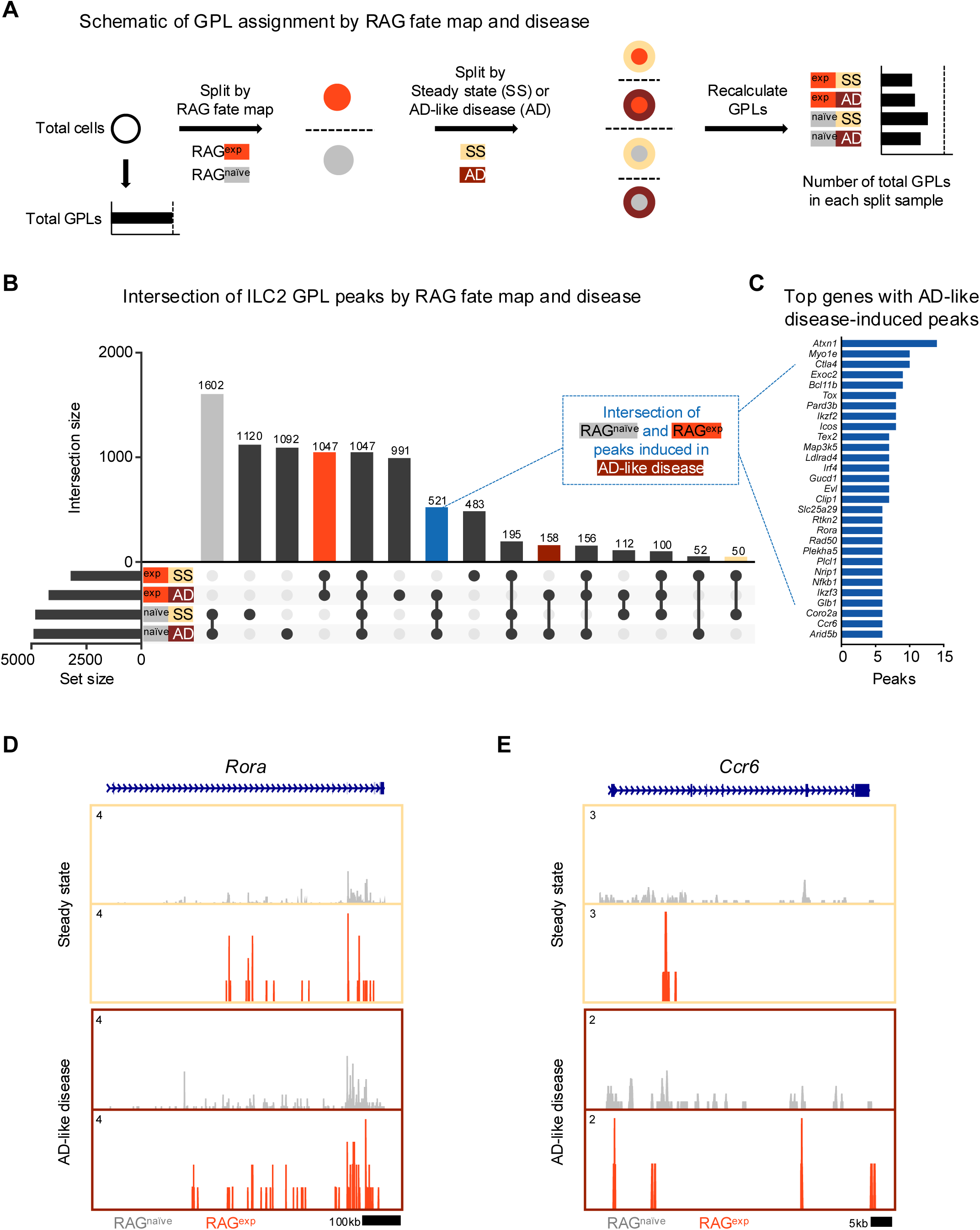
A history of RAG expression broadly influences ILC2 genes at steady state and in AD-like inflammation. **A)** Schematic of the process to determine contribution of RAG fate map and disease states to GPLs for subsequent intersection analyses. GPLs were first calculated for all indictated cells, regardless of disease state or fate map. Cells were then split, first by RAG fate map (RAG^exp^ and RAG^naïve^), and again by disease state (SS - steady state, AD - AD-like inflammation). GPLs were recalculated for each split sample and matched back to the original set of total GPLs. **B)** UpSet plot visualizing intersections of peaks identified from ILC2 GPLs for split samples. Each row represents one of the four sets, and each column corresponds to an intersection of one or more sets (see methods). See **Table S12** for full list of GPLs for all genes. **Table S13** lists total and ILC2 peaks used for intersection analyses in each of the four sets. Columns identifying key intersections are color coded by the corresponding RAG fate map or treatment groups. The blue column indicates the intersection of peaks from RAG^naïve^ cells and peaks induced by AD-like disease in RAG^exp^ cells. **C)** Top genes with the most AD-like disease-induced peaks. Peaks from the intersection between RAG^naïve^ cells and inflamed RAG^exp^ cells were identified in corresponding GPLs, and genes were ranked by number of linked peaks identified. See **Table S14** for full list of ranked genes and associated GPLs. Open chromatin in the ILC2 cell cluster split by disease state and then by RAG fate map for the genomic loci of **C)** *Ccr6* and **D)** *Rora*.

Thus, we next quantified and sorted these GPLs to generate a list of genes with the most peaks "induced" during AD-like disease (**Fig. 6C, Table S14**). Among the identified genes, we selected *Rora* (**Fig. 6C**) and *Ccr6* (**Fig. 6D**) to examine more closely for evidence of epigenomic activation in AD-like disease, given the role of these genes in ILC2 expansion^61,62^ and homing to sites of inflammation^44,104^, respectively. For both genes, we observed more widespread open chromatin over the genomic region in the RAG^naïve^ cells compared to the RAG^exp^ cells, but this difference was partially abolished by increased open chromatin in AD-like disease in the RAG^exp^ cells. Taken together, our analysis reveals that a history of RAG expression selectively modulates activity of ILC2 gene programs across both steady state and during AD-like inflammation, while some programs are more evident at steady state given the uniquely poised nature of ILC2s.

### RAG suppresses the Th2 locus

Our functional data demonstrate a role for RAG expression in regulating ILC2 development and activation, including limiting proportions of IL5^+^ and IL-13^+^ ILC2s at steady state and in AD-like disease. Prior work identified epigenomic priming in ILC2s early in development at the Th2 locus (comprised of the *Il4*, *Il13*, *Rad50*, and *Il5* gene loci) to enable rapid transcriptional responses during inflammation^30^. Thus, we hypothesized that RAG promotes the functional observations in ILC2s by suppressing the establishment of an active regulome at the Th2 locus. To test this hypothesis, we analyzed the Th2 locus in our multiomic data in greater detail. Using a similar strategy to our analysis of ILC2 marker genes, we calculated the number of GPLs in the RAG^exp^ and RAG^naïve^ cells, respectively, for the genes in the Th2 locus. (**Fig. 7A, Table S7**). We found many GPLs associated with the four Th2 locus genes, including significant crosstalk *between* these genes, similar to previous observations (**Fig. 7A**)^105–107^. Importantly, we identified fewer GPLs in the RAG^exp^ cells, particularly for the *Il5* and *Il13* loci (**Fig. 7A**). As in our analysis of the ILC2 marker GPLs, we quantified the differences based on RAG fate mapping and found that all genes in this locus had increased GPLs in RAG^naïve^ cells relative to RAG^exp^ cells (**Fig. 6B; Table S15**). We also applied the same analysis strategy that identified TFs potentially mediating observed differences between RAG^exp^ and RAG^naïve^ ILC2s (**Fig. S10D,E**) specifically to the four Th2 locus genes. Given the limited size of genomic regions (and thus open chromatin peaks) analyzed in the Th2 locus compared to all ILC2 genes, we found overall fewer enriched motifs. Strikingly, significant enrichment of TF motifs was only present in unique peaks from RAG^naïve^ ILC2s, while no TF motifs met the cutoff in RAG^exp^ cells (**Fig S12A, Table S16**). These motifs primarily contained the canonical 5’-(A/T)GATA(A/G)-3’ binding site recognized by the GATA family of zinc finger TFs^108^. When we compared enriched motifs in open chromatin to gene expression of the corresponding TFs, only *Gata3* was expressed at appreciable levels (**Fig S12B**). Critically, *Gata3* expression was higher in RAG^naïve^ compared to RAG^exp^ cells, consistent with our previous analyses of the ILC2 gene regulomes (**Fig. 5C**). Collectively, our data confirm the established role of GATA3 in mediating activation of the Th2 locus^50^ and are consistent with a role for RAG expression in suppressing the type 2 regulome at the Th2 locus.

**Figure 7.**
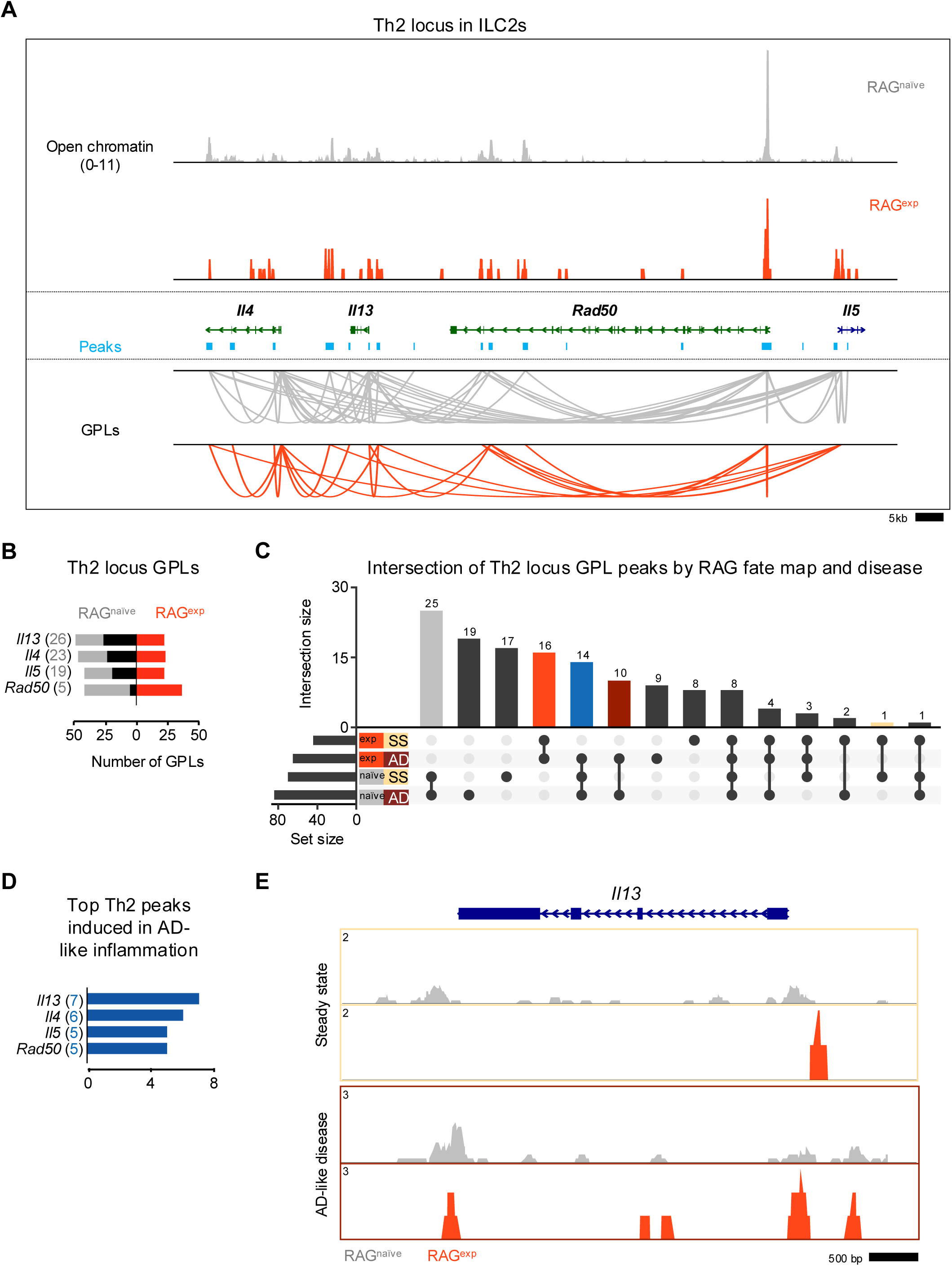
RAG suppresses the Th2 locus. **A)** Coverage plot of the Th2 genomic locus. Open chromatin in the ILC2 cluster for each *Rag1* fate-mapped state is shown on top, and corresponding peaks (teal) and gene to peaks links (GPLs) are shown below for the RAG^naïve^ sample (gray) and the RAG^exp^ sample (tomato red). Only GPLs that fit in the coverage window are shown. **B)** All GPLs identified in each fate map state for the Th2 locus genes *Il4*, *Il13*, *Rad50*, and *Il5*. See **Table S15** for full list of Th2 GPLs. The number of GPLs for each gene is shown on the left in gray for RAG^naïve^ and on the right in tomato red for RAG^exp^. The difference is superimposed in black, and genes are sorted from more GPLs identified in RAG^naïve^ at top to more links identified in RAG^exp^ at bottom. **C)** UpSet plot of intersections of peaks identified from Th2 locus GPLs. GPLs were recalculated, this time for samples separately by both RAG fate map status (RAG^exp^ and RAG^naïve^) and disease (SS - steady state, AD - AD-like inflammation). Each row represents one of the four sets of peaks, and each column corresponds to an intersection of one or more sets (see methods). See **Table S13** for full list of peaks from GPLs for all genes, including Th2 genes, in each of the four sets. Columns identifying key intersections are color coded by the corresponding RAG fate map or disease groups. The blue column indicates the intersection of peaks from RAG^naïve^ cells and peaks induced in AD-like disease in RAG^exp^ cells. (**D)** Th2 genes sorted by number of AD-like disease-induced peaks. Peaks induced by AD-like disease were identified in corresponding GPLs, and genes were ranked by frequency of links to induced peaks (representation in identified GPLs). See **Table S17** for full list of ranked Th2 locus genes and associated GPLs. **E)** Open chromatin tracks, split by disease (beige box – steady state; maroon box – AD-like disease) and by RAG fate map ( RAG^naïve^ - gray, RAG^exp^ - red) for *Il13*.

We next considered the additional effects of AD-like inflammation on the Th2 epigenomic regulome using the same approach we used to analyze the ILC2 gene set in **Fig. 6**. Again, we found the largest set of peaks was shared by the RAG^naïve^ cells, regardless of disease state, with the next largest peak sets belonging to either steady state or AD-like disease in the RAG^naïve^ cells (**Fig. 7C**). Furthermore, there was a large proportion of peaks shared by both RAG^exp^ cells, consistent with a major contribution of a history of RAG expression to epigenomic modulation of the Th2 regulome. To quantify the potential effect of AD-like inflammation on reversing RAG-mediated suppression of Th2 locus genes, we mapped the 14 peaks shared by RAG^naive^ cells and RAG^exp^ cells in the setting of AD-like inflammation (i.e. only suppressed in RAG^naive^ cells at steady state) back to the Th2 genes via their respective GPLs (**Fig. 7C - blue bar, Table S17**). Interestingly, *Il13*, which was not identified as a top ILC2 marker in our earlier analyses, had the highest number of linked peaks associated with potential induction in AD-like disease (**Fig. 7D**). When we examined the *Il13* locus in the ILC2 cluster more closely, we found more widespread open chromatin in the RAG^naïve^ cells compared to the RAG^exp^ cells (**Fig. 7E**). However, in the AD-like disease sample, the RAG^exp^ cells displayed increased open chromatin relative to the steady state, consistent with induction in the setting of inflammation, like our earlier findings for ILC2 genes such as *Ccr6* (**Fig. 6E**). Taken together, our functional data and multiomic analyses demonstrate a role for RAG expression in modulating genes critical for ILC2 development and function, including the key type 2 cytokines expressed from the Th2 locus.

## DISCUSSION

RAG recombinases evolved nearly 500 million years ago from endogenous transposons, crucially enabling antigen receptor rearrangement and emergence of the adaptive immune cell lineages present in all modern vertebrates^29,109^. Indeed, RAG deficiency leads to a complete lack of B and T lymphocytes, manifesting clinically as severe combined immunodeficiency (SCID)^110–112^. However, fate mapping studies have shown that multiple mature immune cell populations other than adaptive B and T lymphocytes have a history of RAG expression^23,24,33,113,114^. More recent studies by Karo et al found that RAG expression during NK cell development influences multiple cellular functions including antitumor cytotoxicity, cell proliferation, and survival^24^. Yet whether RAG modulates cellular functions of innate immune cell populations other than NK cells remains poorly understood. Here, using RAG-deficient mice, RAG fate mapping mice, and multiomic analyses, we report that RAG suppresses developmental and effector functions of ILC2s.

Our functional data in RAG-deficient mice demonstrate that populations of ILC2s producing the type 2 cytokines IL-5 and IL-13 preferentially expand in the absence of a history of RAG expression. This implies a specific role for RAG in developmental repression of ILC2s. Building on this, our multiomic RAG fate mapping analyses of ILC2 gene programs demonstrate extensive epigenomic differences between RAG^exp^ and RAG^naïve^ cells. We found RAG-associated epigenomic suppression at multiple functional levels, including cell surface receptors, key transcription factors, and the Th2 locus encoding the type 2 cytokine genes *Il5* and *Il13*. Although RAG is only transiently expressed early in lymphoid development^115^, our data demonstrate that RAG expression can imprint durable effects on ILC2 gene programs to restrain their function.

Our observations imply that RAG expression may mark a developmentally distinct population of ILC2s. In adaptive lymphocytes, RAG expression in T cells is restricted to their time in the thymus. However, ILC2 populations have been observed in the thymus, provoking the hypothesis that thymic ILC2s may be uniquely high in expression of RAG^63,116^. Prior studies by Schneider et al have identified ILC2 populations in adult tissues that variably derive from expansion of fetal, postnatal, and adult populations^68^. Yet how RAG expression in ILC2s may be restricted spatially or temporally remains unknown. The mouse strains used in the fate mapping studies by Schneider et al would be incompatible with our RAG fate mapping mice. Thus, novel mouse strains enabling intersectional genetics to trace ILC2 ontogeny (e.g. CreER/lox for temporally restricted fate mapping and FlpO/frt for RAG fate mapping^117^) are needed to more precisely determine when and where RAG expression occurs during ILC2 development. Beyond steady state ontogeny, our data suggest a history of RAG expression also imprints suppressed proliferative and type 2 inflammatory functions on ILC2s in the setting of AD-like disease.

It is increasingly recognized that expression of effector molecules for both ILCs and their counterpart adaptive lymphocytes (e.g. IL-13 from ILC2s and Th2 cells) is governed by finely tuned transcriptomic and epigenomic regulomes^30,118–120^. ILCs tend to adopt these regulomes earlier in their development than T cells, and these “poised” regulatory elements are thought to underlie the ability of tissue-resident ILCs to rapidly respond to stimuli. In contrast, the regulomes of naïve T cells remain relatively inactive until stimulation. Given that T cells are uniformly RAG-experienced, our data provoke the hypothesis that RAG^exp^ ILC2s adopt a phenotype closer to that of naive T cells and may require stronger stimuli than RAG^naïve^ ILC2s to become activated. Indeed, our analyses found the RAG-associated suppressive programs could be overcome in the setting of AD-like inflammation. Thus, sufficient RAG expression may mediate key events underlying the establishment and maintenance of functional regulomes not only in ILCs, but also T cells. How RAG might affect these changes, and whether they are independent of its enzymatic activity and/or antigen receptor recombination, remains to be elucidated.

Clinically, a link between enhanced type 2 immune activity and RAG dysfunction is well-established. Omenn Syndrome (OS) is a form of SCID characterized by exaggerated type 2 immune activation and typically arises in the setting of hypomorphic RAG gene mutations. Impaired antigen receptor rearrangement, with rare “leaky” recombination events, leads to expansion of autoreactive oligoclonal T cells, eosinophilia, and markedly elevated IgE^121–125^. Similar phenotypes have been observed in mice harboring RAG mutations analogous to those found in human patients with OS^126,127^. Notwithstanding these findings, the mechanisms underlying the propensity of oligoclonal T cells with hypomorphic RAG activity to preferentially develop into the Th2 subtype are unclear. Prior studies have found a role for regulatory T cells in controlling type 2 skewing of transferred T cells in RAG-deficient hosts, potentially explaining similar observations in patients with OS^128^. Our data provide an additional mechanism by which RAG dysfunction may lead to OS through loss of cell-intrinsic RAG-mediated suppression of type 2 cellular programs. Additionally, increased type 2 cytokine production from RAG-deficient ILC2s may, in *trans*, enhance expansion of the oligoclonal Th2 cell populations, IgE induction, and eosinophilia observed in RAG-deficient states like OS. However, whether other immune cell types with RAG dysfunction, such as ILCs, contribute to the pathogenesis of OS in humans has not been investigated.

Lymphoid acquisition of RAG activity may represent a newer evolutionary mechanism that fine tunes ancient innate immune cell programs in addition to enabling development of relatively newer antigen-specific adaptive immune cell populations. Independent of antigen receptor diversity, loss of this function may offer an explanation as to why oligoclonal T cells tend to expand and skew towards a Th2 cell phenotype in the setting of hypomorphic RAG function as in OS^128^. Further studies are needed to define whether the suppressive effects of RAG expression operate similarly in T and B cells. Although we demonstrate that this phenomenon is observed in ILC2s, whether hypomorphic RAG expression in bona fide Th2 cells not only results in oligoclonality but also loss of suppression of the Th2 locus independently of antigen receptor rearrangement remains an outstanding question. Indeed, during development of gene therapy strategies for RAG-deficient SCID, lower doses of wild type RAG transgene expression have been associated with development of OS-like conditions in transplanted RAG-deficient mice^129–132^.

A major limitation of our study is a focus on cutaneous type 2 inflammation, which stemmed from our initial observations in the MC903 mouse model of AD-like disease. Further, given the scarcity of skin-resident ILC2 populations, key functional investigations in our study such as cytokine production and multiomic sequencing were limited to the sdLN, as in prior studies^13,133^. However, ILC2s are recognized to have highly tissue-specific functions that extend much beyond inflammation to other processes including regeneration and metabolism. In addition to Il-5 and IL-13, ILC2s can produce other effector molecules such as acetylcholine, IL-9, methionine-enkephalin peptides, and amphiregulin, which modulate tissue responses across numerous organs^134–141^. Considering that the complexity of ILC2 biology may result in markedly divergent responses to RAG expression in other tissues and disease models, we thus restricted our initial studies to the skin, where we had strong molecular, cellular, and phenotypic outcomes. An implication of our findings in the skin is that RAG expression may modulate a variety of ILC2 functions in other tissues. Broader surveys of how RAG impacts ILC2 development and function in different tissues and disease states remains an exciting area of inquiry.

While we focused our multiomic analyses on ILC2s, it is likely that RAG may impact other ILC populations. For example, hyperactivation of intestinal ILC3s has been observed in *Rag1^-/-^* mice secondary to persistent phosphorylation of Signal Transducer and Activator of Transcription 3 (STAT3). Adoptive transfer of T regulatory cells rescued this phenotype, providing a cell-extrinsic mechanism for the observation of hyperactivated ILC3s in the setting of RAG deficiency^142^. However, our data supporting a cell-intrinsic role for RAG in ILC2s may offer additional mechanistic insight into the prior observations in ILC3s. We found that the regulome of *Jak2*, which encodes JAK2, an upstream activator of STAT3, was more activated in RAG^naïve^ ILC2s (**Fig. 5C**). Additionally, the regulome for *Dusp1*, which encodes dual specificity phosphatase 1 (DUSP1), was more activated in RAG^exp^ ILC2s (**Fig. 5C**). While not implicated in directly dephosphorylating STAT3, a recent study found DUSP1 overexpression negatively regulated the JAK2/STAT3 pathway^143^. Notably, recent transcriptional profiling of skin ILCs identified a potential mechanism for skin ILC populations to transition to an ILC3-like phenotype^69^, but how this process is regulated remains poorly understood. Taken together, our data provoke compelling new hypotheses about cell-intrinsic functions of RAG that may be complementary, rather than contradictory, to prior observations in gut and skin ILC populations. Additionally, our studies provide a rationale to design novel reagents to enable more comprehensive studies on the role of RAG in multiple innate immune cell populations across different tissues and disease models.

Our observations are also limited by lack of a defined mechanism for how RAG expression imprints durable epigenomic and transcriptomic changes in ILC2s. The mechanisms by which RAG mediates VDJ recombination are well-defined, from the biochemical details of DNA-binding to the epigenomic accessibility of antigen receptor loci and timing of RAG expression^29,144–146^. Notwithstanding genomic stress^24^ or potential RAG dose effects^129–132^, how RAG expression might modulate broad developmental and functional lymphoid programs other than V(D)J recombination remains unclear. The RAG complex can bind both DNA and modified histones and has been observed to occupy thousands of sites across the genome^27^. Thus, RAG may directly influence open chromatin states or obscure transcription factor binding sites to alter ILC2 development and function. Notably, RAG preferentially binds near transcription start sites of open chromatin in mouse thymocytes and pre-B cells, although corresponding effects on gene expression were not observed^27^. Although canonical recombination sites are concentrated in the antigen receptor loci, cryptic recombination sites in other regions may be deleted or rearranged by RAG activity, altering transcriptional regulation of associated genes^27^. In contrast to developing B and T lymphocytes, the precise timing and location of RAG expression in ILC2s is not known. Thus, combined with the relative scarcity of ILC2s, conventional methods of chromatin immunoprecipitation to identify potential epigenomic regulatory mechanisms mediated by RAG expression may not be feasible in ILC2s or other rare cell populations. Instead, newer technologies such as self-reporting transposons^147^ could be adapted to trace the genomic footprint of RAG in cells at various stages of development and in various tissues independent of the constraint of concurrent RAG expression. Finally, through its E3 ubiquitin ligase activity^29^, RAG may influence immune signaling pathways independently of transcription altogether. Given that direct targeting of RAG would lead to unacceptable side effects, elucidating the mechanisms by which RAG imprints phenotypic changes beyond antigen receptor rearrangement is a critical next step in translating these findings to potential new therapies.

Our studies expand prior work implicating RAG in critical immune functions beyond antigen receptor rearrangement that is exclusive to adaptive lymphocytes. Further, we provide additional insights into why patients with OS exhibit atopic syndromes in the setting of adaptive lymphocyte deficiency. Future studies into mechanisms underlying these findings may lead to new therapeutic avenues for disorders such as atopic dermatitis, food allergy, and asthma.

## Supporting information

zip of all supplemental files

## ACKNOWLEDGEMENTS

We thank all members of the Kim lab for helpful comments and discussion. This work is supported by the Allen Discovery Center program, a Paul G. Allen Frontiers Group advised program of the Paul G. Allen Family Foundation, the Doris Duke Charitable Foundation, LEO Pharma, the National Institute of Arthritis and Musculoskeletal and Skin Diseases (NIAMS) (AR070116, AR077007, and AR080392), and the National Institute of Allergy and Infectious Disease (NIAID) (AI167933 and AI167047) of the National Institutes of Health (NIH). A.M.V. is supported by NIAMS (1K08AR080219). M.T. is supported by the Japanese Society of Allergology (JSA) International Scholarship. A.M.T. is supported by the NIAID (AI007163 and AI154912). We thank the Genome Technology Access Center at the McDonnell Genome Institute at Washington University School of Medicine for help with genomic analysis. The Center is partially supported by NCI Cancer Center Support Grant #P30 CA91842 to the Siteman Cancer Center and by ICTS/CTSA Grant# UL1TR002345 from the National Center for Research Resources (NCRR), a component of the National Institutes of Health (NIH), and NIH Roadmap for Medical Research. This publication is solely the responsibility of the authors and does not necessarily represent the official view of NCRR or NIH.

## AUTHOR CONTRIBUTIONS

Conceptualization: A.M.V., M.M., L.Z., M.T., T-L.Y., and B.S.K.

Methodology: A.M.V., M.M., M.T., L.Z., T-L.Y., and B.S.K.

Validation: A.M.V., M.M., L.Z., M.T., and T-L.Y.

Formal Analysis: A.M.V. and M.M.

Investigation: A.M.V., M.M., L.Z., M.T., T-L.Y., D-H.K., and H.J-M.

Resources: S.V.D. and B.S.K.

Writing – Original Draft: A.M.V.

Writing – Review & Editing: All authors

Supervision, B.S.K.

Funding Acquisition, B.S.K.

---

**Figure S1.**
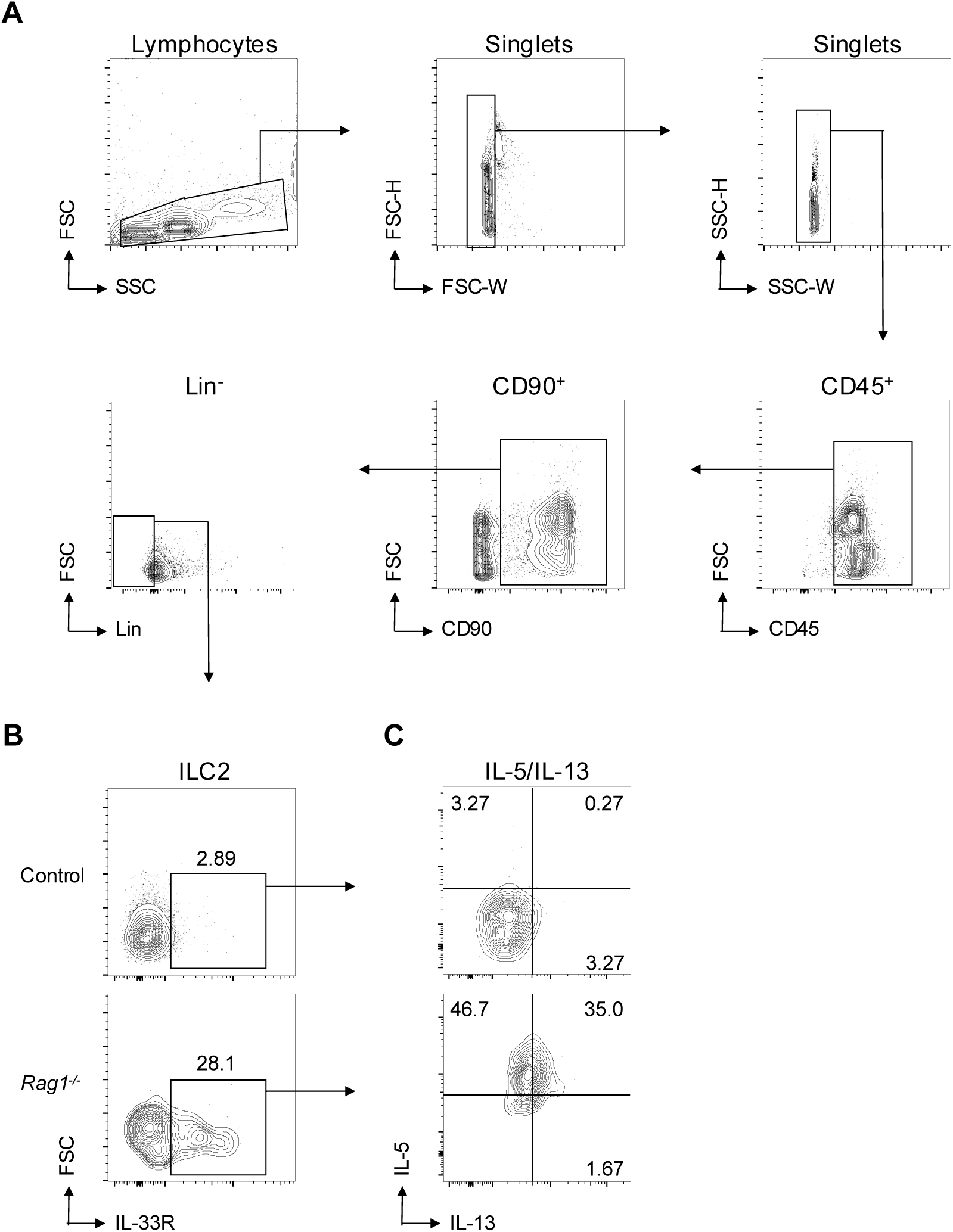
ILC2 and IL-5/IL-13 gating. Gating for **A)** CD45^+^, CD90^+^, Lin^-^ cells (Lin^-^ defined as CD3^-^, CD5^-^, CD11b^-^, CD11c^-^, CD19^-^, NK1.1^-^, and FcεR1^-^), then gating on **B)** ILC2 (IL-33R^+^ Lin^-^) corresponding to Fig. 1C, with subsequent gating of **C)** IL-5^+^ and IL-13^+^ ILC2, corresponding to Fig. 1D**-E**.

**Figure S2.**
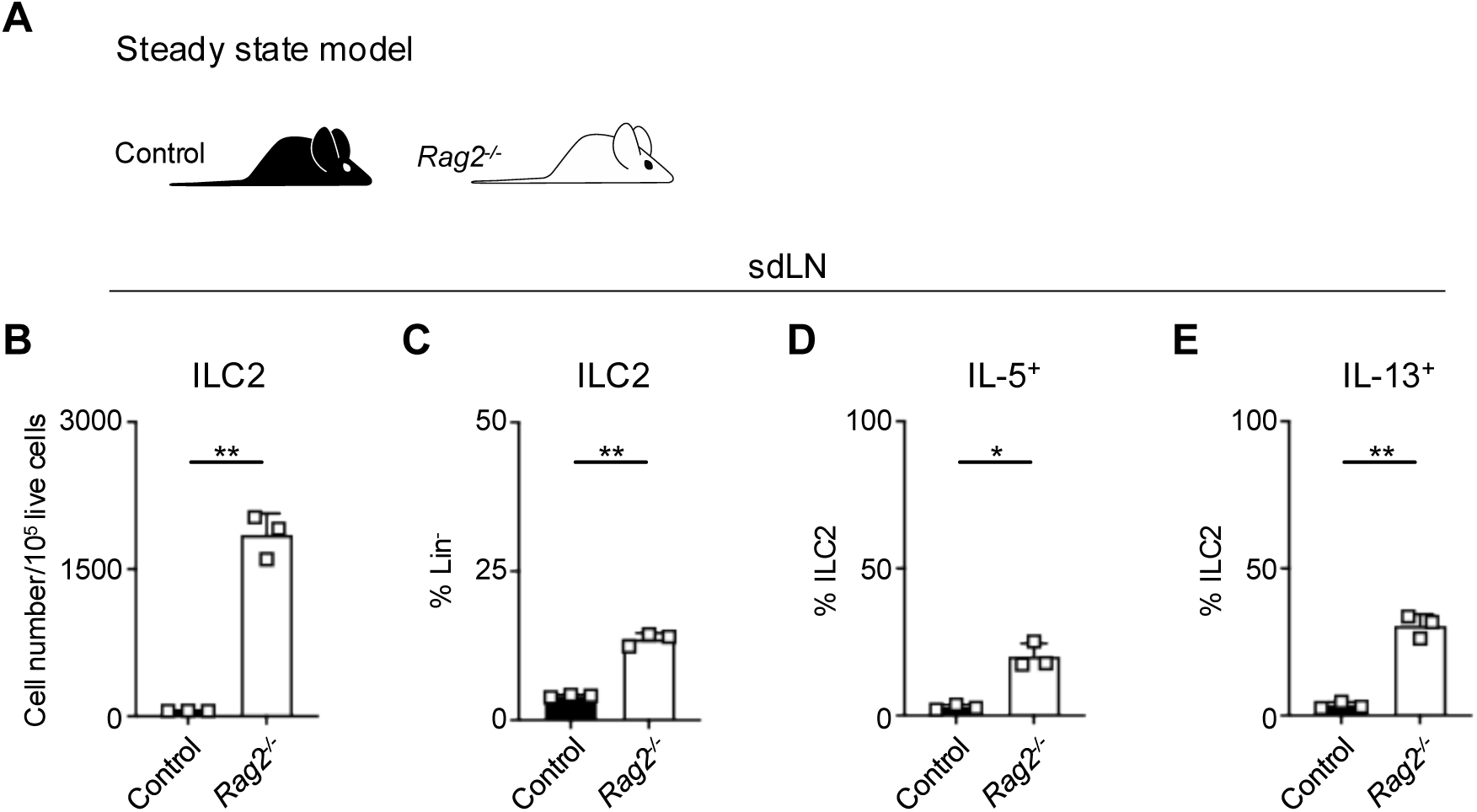
Expansion and activation of ILC2s in RAG2 deficiency compared to littermates. **A)** Schematic of steady state analysis of WT B6 (Control) mice or *Rag2^-/-^*mice. **B)** Proportion of CD90^+^ Lin^-^ cells (Lin-defined as CD3^-^, CD5^-^, CD11b^-^, CD11c^-^, CD19^-^, NK1.1^-^, and FcεR1^-^) determined to be ILC2s (IL-33R^+^) in sdLN at steady state from WT or *Rag2^-/-^* mice. Percent ILC2 from sdLN at steady state following PMA/iono stimulation positive for **C)** IL-5 or **D)** IL-13 staining. Data representative of 2 independent experiments, 2-3 mice per group. * P <0.05, ** P < 0.01 by two-tailed Welch’s t test. All data represented as mean with standard deviation.

**Figure S3.**
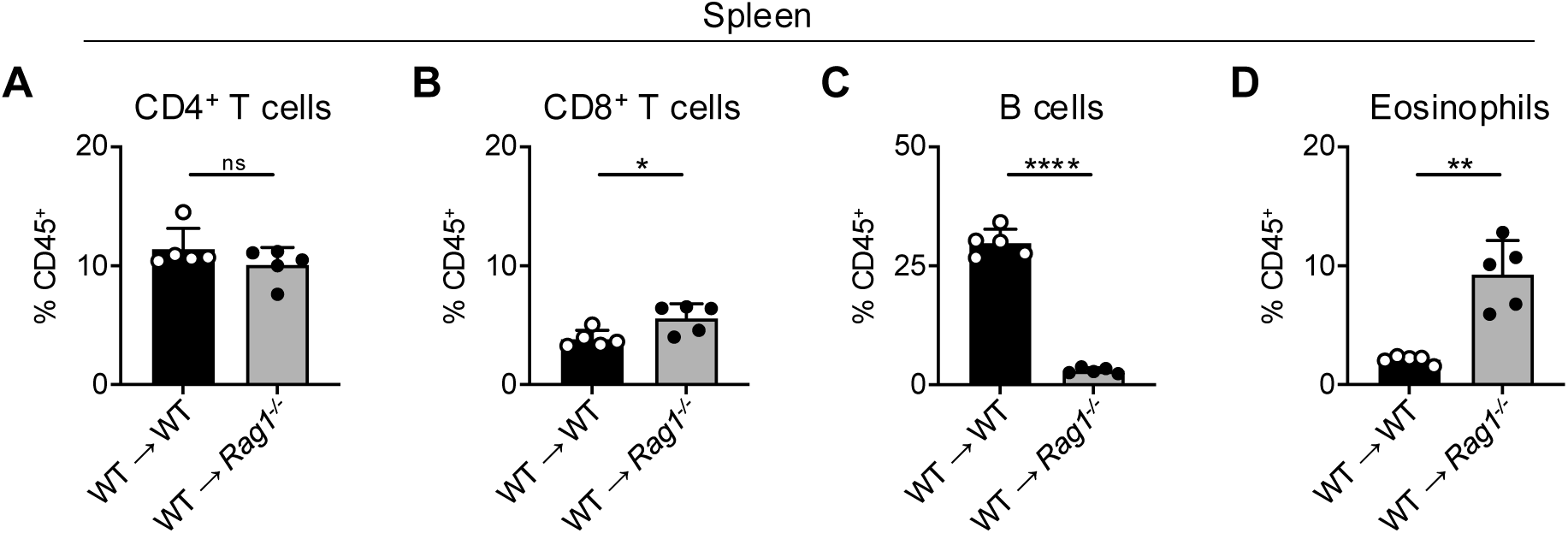
Confirmation of splenocyte reconstitution in splenocyte chimera mice. Proportion of CD45^+^ splenocytes from splenocyte chimera mice (related to Fig. 2A**-E**) determined to be **A)** CD4^+^ T cells (CD4^+^, CD8^-^, CD19^-^), **B)** CD8^+^ T cells (CD4^-^, CD8^+^, CD19^-^), **C)** B cells (CD4^-^, CD8^-^, CD19^+^), and **D)** Eosinophils (SiglecF^+^, CD4^-^, CD8^-^).

**Figure S4.**
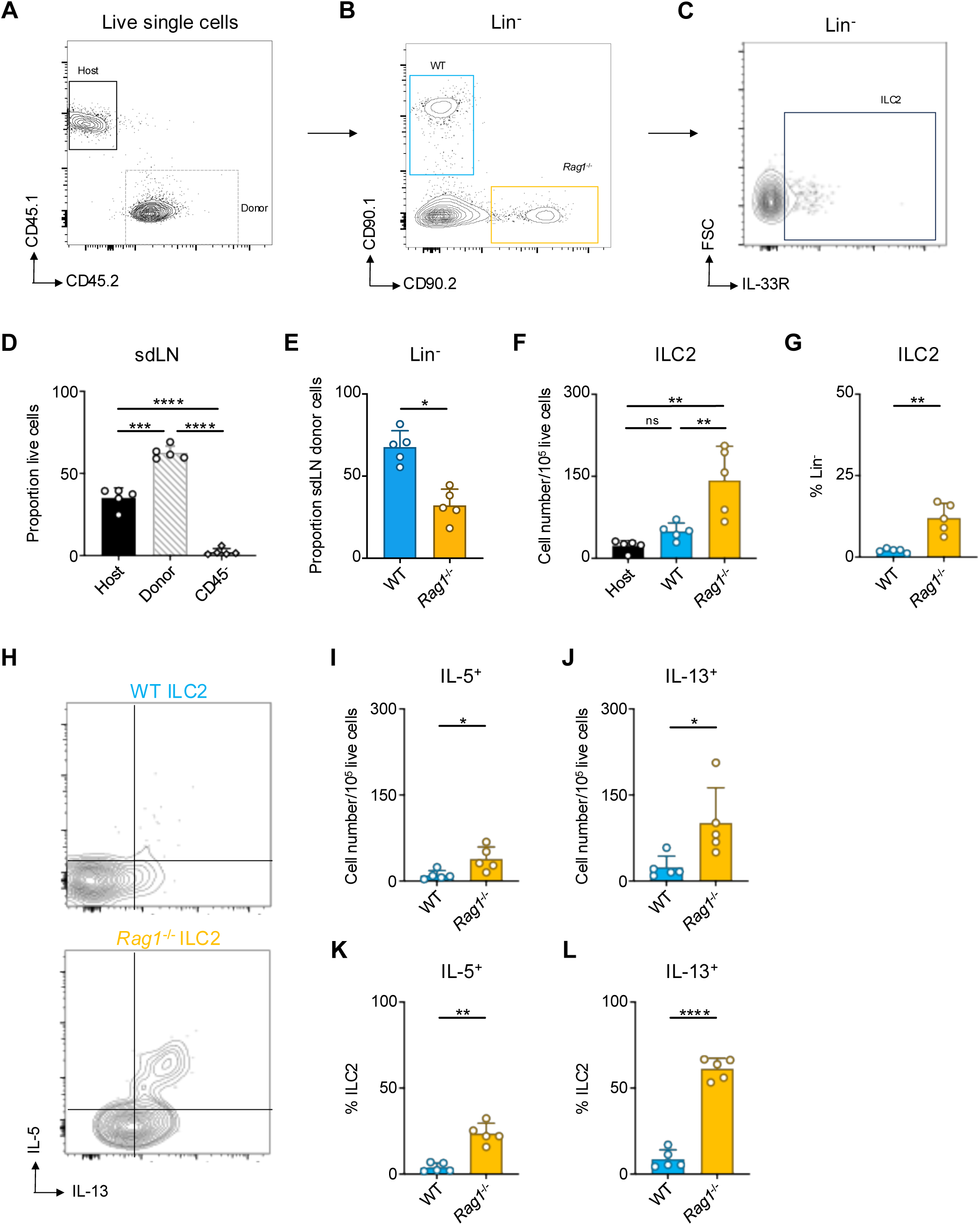
Donor cell reconstitution and gating in sdLN of WT:*Rag1^-/-^* bone marrow chimera mice. **A)** Gating of live host (CD45.1^+^) and donor (CD45.2^+^) cells then **B)** Gating on donor cells by genotype (CD90.1^+^ = WT [blue]; CD90.2^+^ = Rag1^-/-^ [orange]) in Lin^-^ population then **C**) Gating on Lin^-^ IL-33R^+^ ILC2s in the skin draining lymph node (sdLN). **D**) Host/donor CD45^+^ cell reconstitution in sdLN of WT:*Rag1^-/-^* bone marrow chimera mice. **E)** quantification of CD45.2^+^ Lin^-^ donor cells by genotype in sdLN of WT:*Rag1^-/-^*bone marrow chimera mice. **F**) total numbers of ILC2s normalized to 10^5^ live cells and **G**) ILC2 proportion of Lin-cells in the sdLN. **H**) Gating for IL-5 and IL-13 after in vitro stimulation and intracellular cytokine staining of ILC2s from sdLN. Quantification of total positive cells normalized to 10^5^ live cells for **I**) IL-5 and **J**) IL-13 and proportion of ILC2 positive for **K**) IL-5 and **L**) IL-13. **E,G,I-L:** * P < 0.05, ** P <0.01 by ratio means paired t test. **D,F**: ** P <0.01, *** P < 0.001, **** P < 0.0001, by RM one-way ANOVA test with Geisser-Greenhouse correction. All data represented as mean with standard deviation. Related to Fig. 2G**-J**.

**Figure S5.**
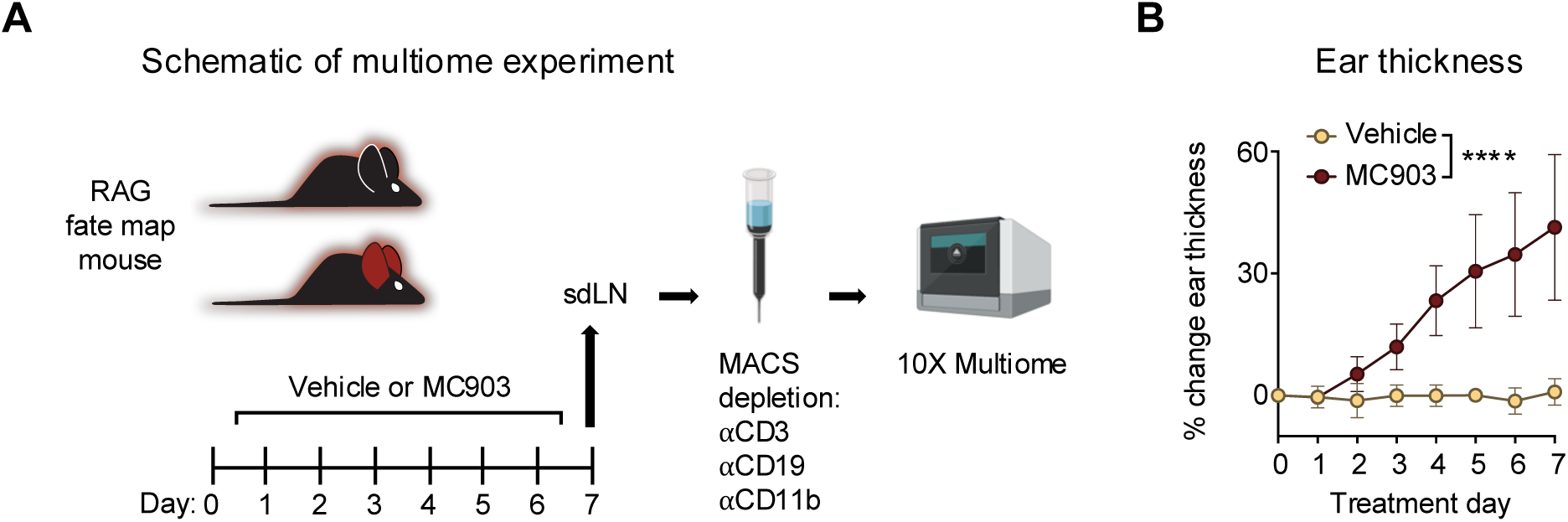
sdLN multiome experiment. **A)** *Rag1*^Cre^::*Rosa26*^LSL-tdRFP^ reporter mice were given topical treatments with 2 nmol MC903 dissolved in ethanol vehicle or with ethanol vehicle alone to each ear daily for 7 days. Harvested sdLN processed using Magnetic Activated Cell Sorting (MACS) led to depletion of cells expressing the CD3, CD19, and CD11b lineage markers and remaining cells were further processed in the 10X Multiome pipeline, generating both single cell RNA-sequencing and single cell ATAC-sequencing data for each cell. **B)** Ear thickness measured daily in the AD-like disease multiome experiment. Data representative of one experiment, with 4 mice per group pooled for sequencing. **** P <0.0001 by 2-way ANOVA with Sidak’s multiple comparisons test, day 7. All data represented as mean with standard deviation.

**Figure S6.**
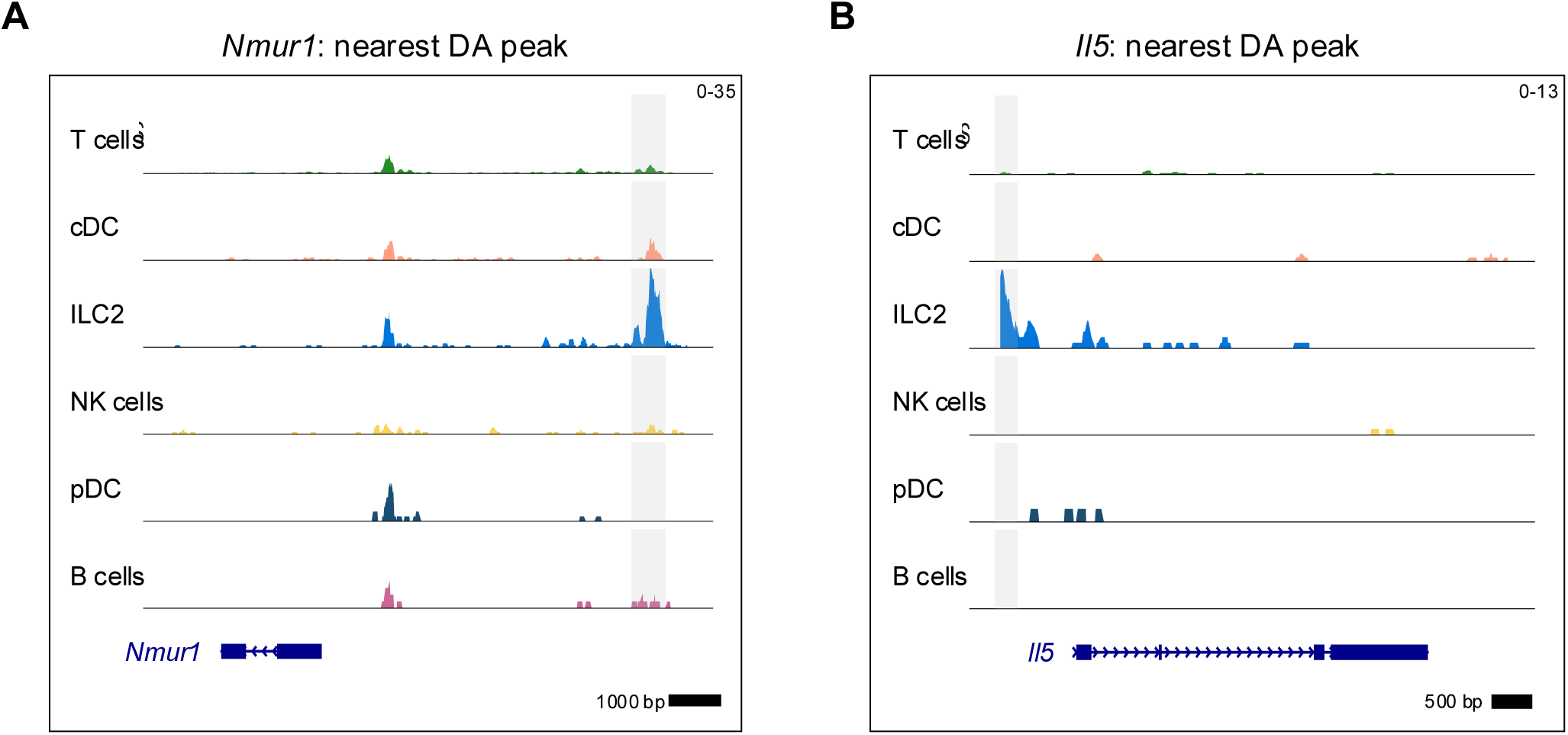
ILC2 marker genes identified in the differentially accessible open chromatin assay. Differentially accessible (DA) open chromatin peaks identified for the ILC2 cluster are highlighted in gray and shown next to the closest gene for **A)** Neuromedin U receptor 1 (*Nmur1*) and **B)** IL-5 (*Il5*). See **Table S3** for top 100 DA peaks and distances to nearest genes for the ILC2 cluster.

**Figure S7.**
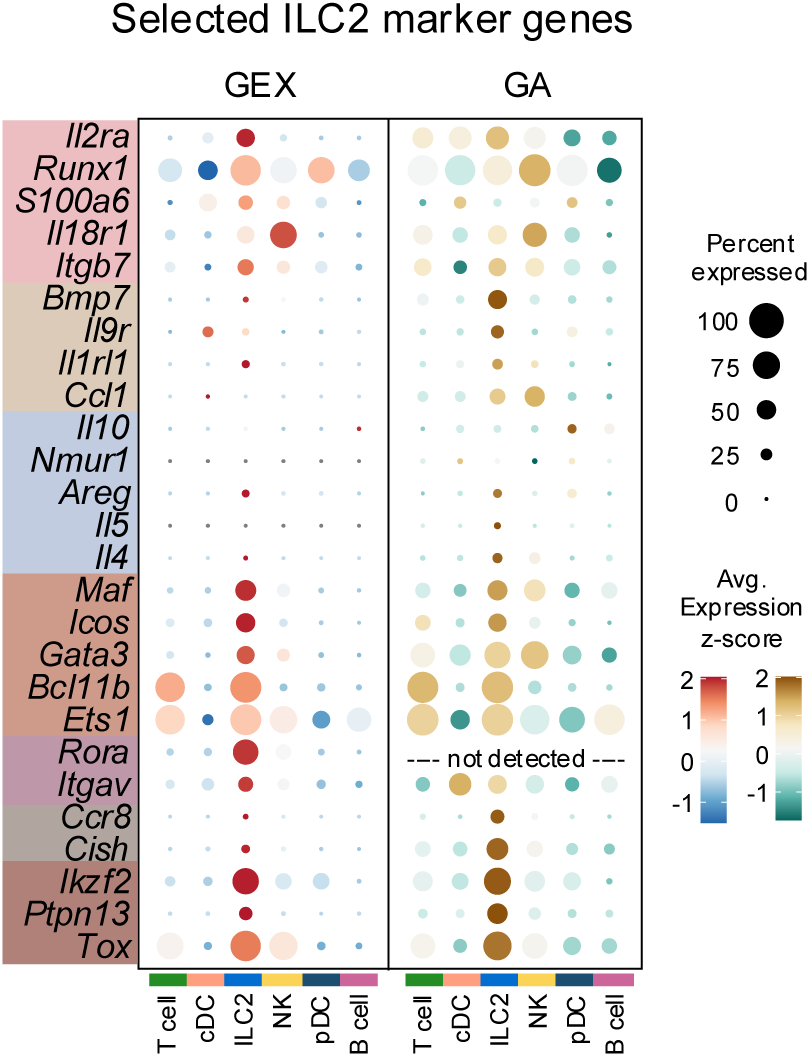
Dotplots of selected ILC2 marker genes. Dotplots comparing selected marker genes from the multiomic ILC2 gene set (Figure 4I**, Table S4**) for each cluster between the GEX and GA assays, with genes highlighted by color corresponding to individual assays or overlap of assays in which they were identified. *Rora* was not detected in the GA assay.

**Figure S8.**
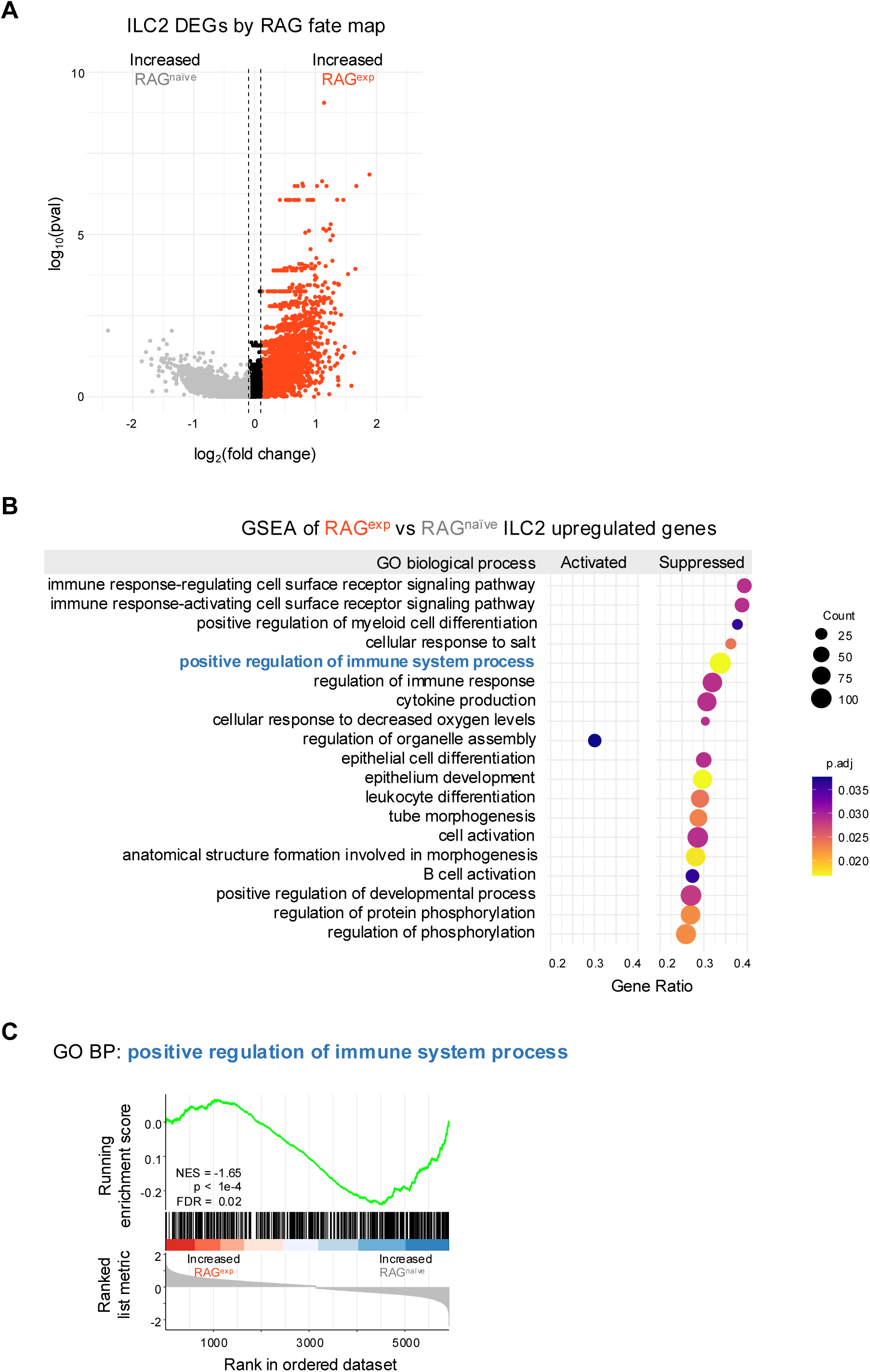
Gene set enrichment analysis of differentially expressed genes in ILC2s. **A)** Volcano plot of differentially expressed genes (DEGs) by RAG fate map for the ILC2 cluster. A ranked list (**Table S5**) was constructed for all DEGs with log_2_(fold change) >0.1 for gene set enrichment analysis (GSEA). **B)** Dotplot of GSEA result calculated using ClusterProfiler and the gene ontology (GO) biological process (BP) database (see methods). Full results in **Table S6**. **C)** GSEA plot of the GO BP “positive regulation of immune system process” gene set. RAG^naïve^, RAG fate map negative; RAG^exp^, RAG fate map positive.

**Figure S9.**
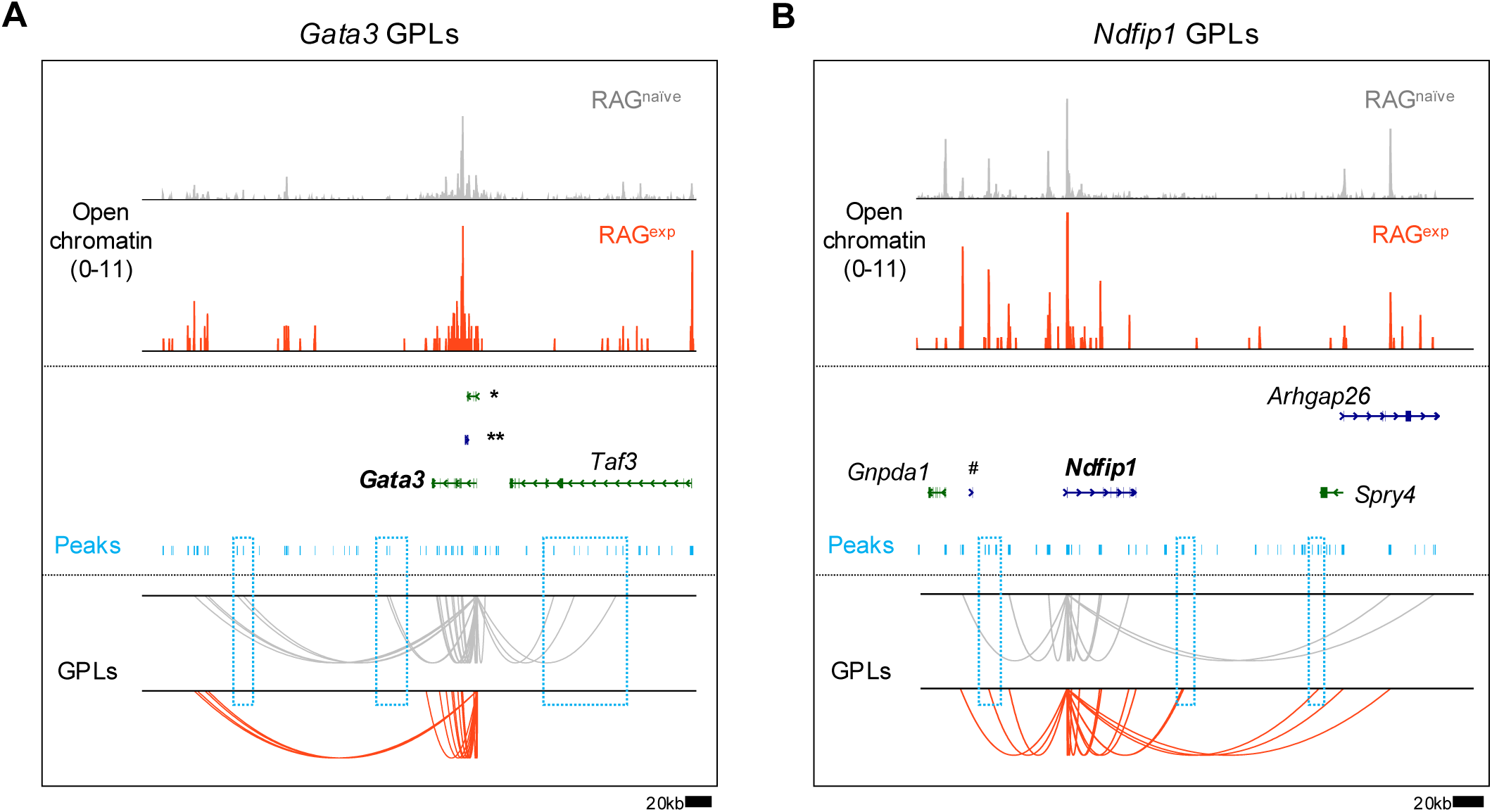
Mapping gene to peak links in select ILC2 genes. Gene to peak links (GPLs) mapped for the RAG^naïve^ and RAG^exp^ states as depicted in Figure 5B for **A)** GATA binding protein 3 (*Gata3*) and **B)** Nedd4 family interacting protein 1 (*Ndfip1*). Only GPLs that fit in the coverage window are shown. Select peaks (teal bars) present in one state, but not the other, are highlighted in teal boxes. Full gene names not shown on figure in (A) are \**9230102O04Rik* and \*\**4930412O13Rik* and in (B) ^#^*Gm42690*.

**Figure S10.**
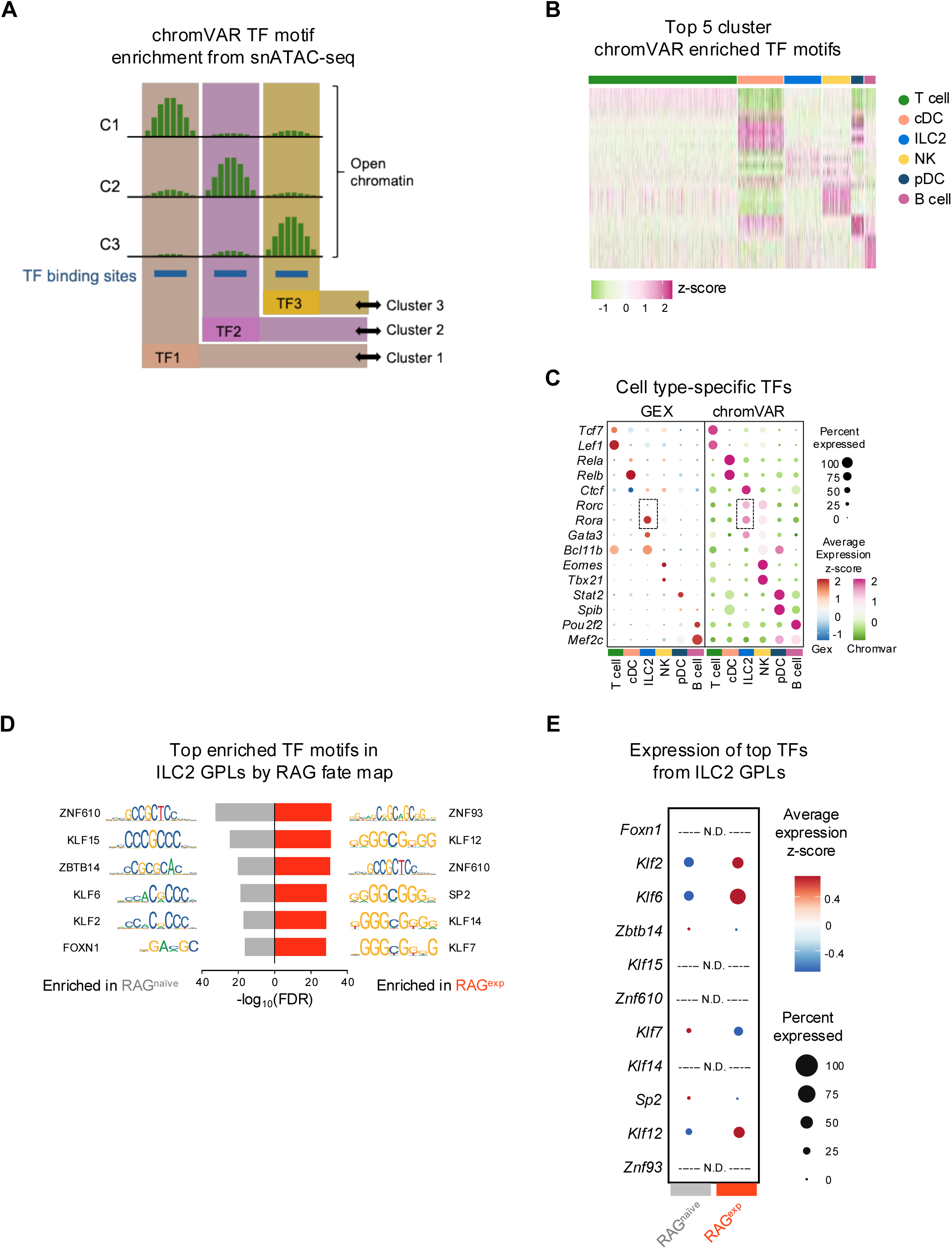
Multiomic transcription factor analysis of ILC2s. **A**) Schematic of assigning transcription factor (TF) motif enrichment in differentially open chromatin to cell clusters using chromVAR. **B**) Heatmap of top 5 TF motif activity scores for each cluster. **C**) Dotplots comparing expression levels of selected TFs in the gene expression (GEX) assay with the chromVAR activity score of the corresponding TF motif. The TFs and corresponding motifs for *Rora* and *Rorg* in the ILC2 cluster are highlighted by boxes. An expanded list of cluster TF motif markers identified using chromVAR is in **Table S10**. **D**) Analysis of TF motifs enriched in ILC2 gene to peak links (GPLs) unique to RAG^naïve^ and RAG^exp^ populations determined using the FindMotifs function in Signac. The top 6 TF motifs for each population are shown and are ranked by the −log_10_ transformed false discovery rate (FDR - Bonferroni corrected p values). An expanded list is in **Table S11**. **E**) Dotplot of gene expression for TFs corresponding to the top enriched TF motifs identified in ILC2 GPLs from (D). TF genes that were not detected in the GEX assay are labeled N.D.

**Figure S11.**
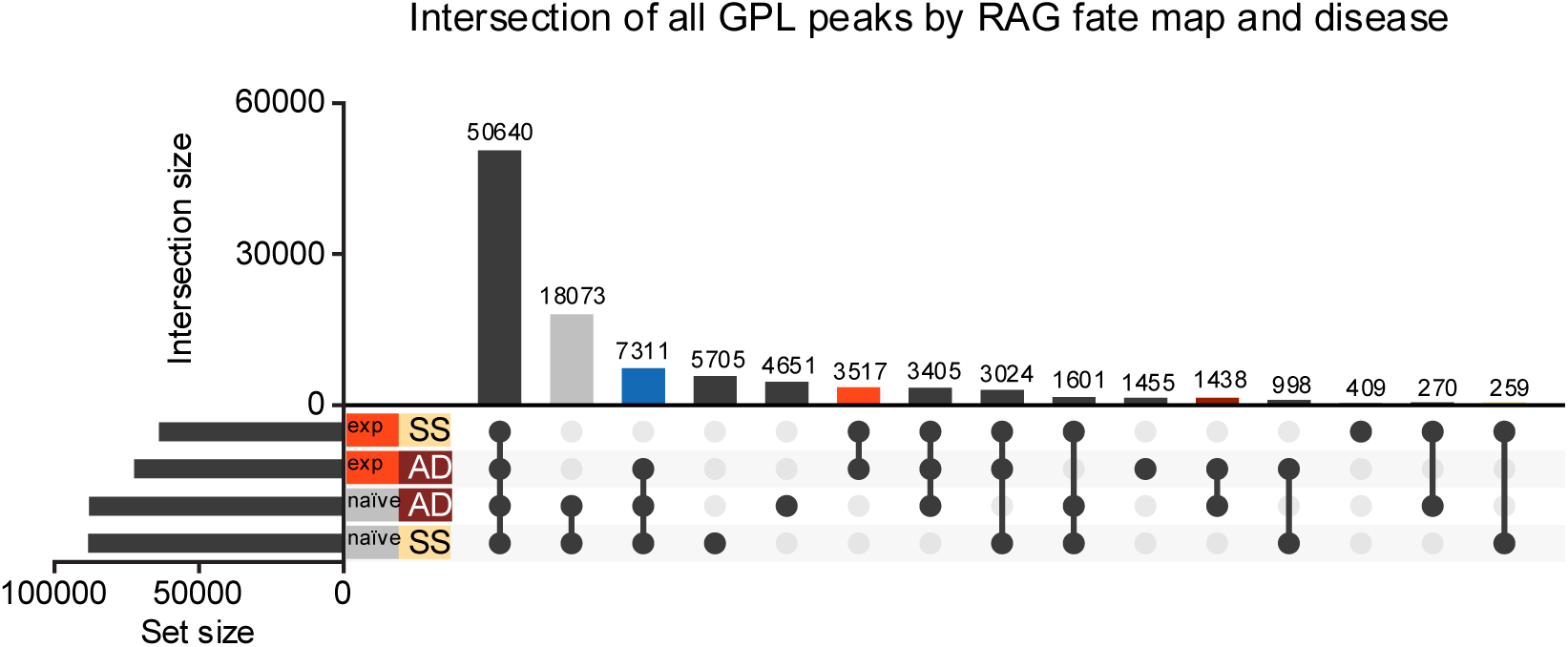
Gene to peak link analysis by RAG fate map and disease for all detected genes. **A)** UpSet plot of overlaps in peaks identified from GPLs of all genes split by both RAG fate map (RAG^exp^ and RAG^naïve^) and disease state (SS - steady state, AD - AD-like inflammation). Each row corresponds to one of the four sets, and each column corresponds to an intersection of one or more sets (see methods). See **Table S13** for full list of peaks from GPLs for all genes in each set. Columns identifying key intersections are color coded by the corresponding RAG fate map or disease groups. The blue column indicates the intersection of peaks from RAG^naïve^ cells and peaks induced by AD-like disease in RAG^exp^ cells.

**Figure S12.**
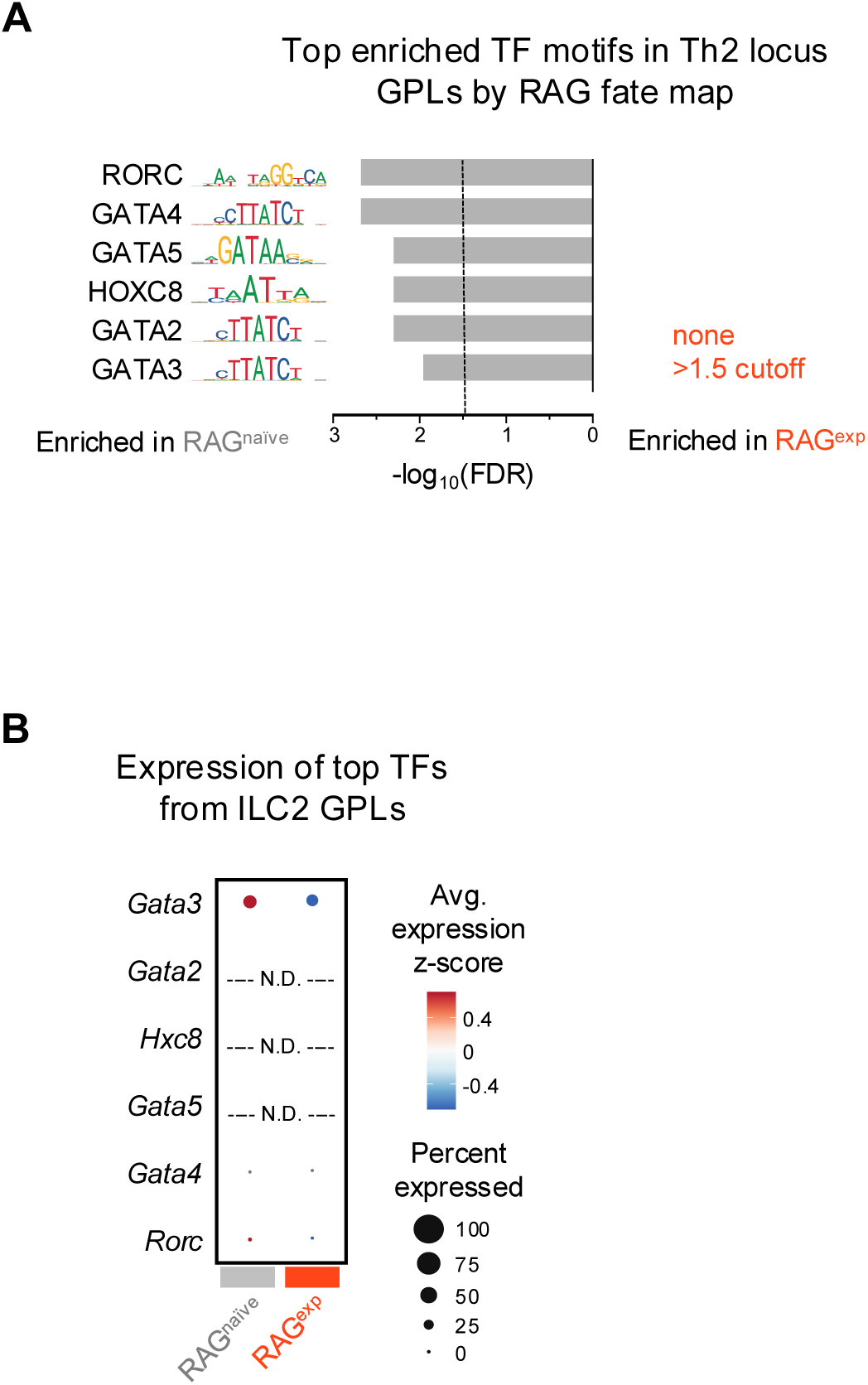
Multiomic transcription factor analysis of Th2 locus. **A**) Analysis of TF motifs enriched in Th2 locus subset of ILC2 gene to peak links (GPLs) unique to RAG^naïve^ and RAG^exp^ cell populations determined using the FindMotifs function in Signac. The top 6 TF motifs for each population are shown and are ranked by the −log_10_ transformed false discovery rate (FDR - Bonferroni corrected p values). No TFs met the −log10(FDR) minimum cutoff value of 1.5 in the RAG^exp^ cell population. The full list of enriched motifs is in **Table S16**. **B**) Dotplot of gene expression for TFs corresponding to the top enriched TF motifs identified in Th2 locus GPLs from (A). TF genes that were not detected in the GEX assay are labeled N.D.

## MATERIALS AND METHODS

### Animal studies

Wild-type (WT) C57BL/6J and WT congenic strains (CD90.1^+^, CD45.1^+^), *Rag1*^-/-^, and *Rag2*^-/-^ mice were initially purchased from the Jackson Laboratory and bred in house. The RAG fate-mapping strain *Rag1*^Cre^::*Rosa26*^LSL-tdRFP^ was originally created in the lab of Paul Kincade^33^ and bred in house. All mice were housed in specific-pathogen-free condition and environmentally controlled animal faculty with a 12-hour light-dark cycle and given unrestricted access to food and water at Icahn School of Medicine at Mount Sinai or Washington University School of Medicine in St. Louis. All animal protocols and experiments were approved by the Institutional Animal Care and Use Committee (IACUC) at Icahn School of Medicine at Mount Sinai or Washington University School of Medicine in St. Louis. Experiments were performed on independent cohorts of male and female mice. For induction of AD-like disease, 8- to 12-week-old mice were treated with 2 nmol calcipotriol (MC903, Tocris Bioscience) in 10 μL of 100% ethanol (EtOH) vehicle, or vehicle alone, on the bilateral ear skin daily for 7-10 days. Body weight and ear thickness were measured daily with a digital scale and analog caliper by the same investigator. For tissue harvest, animals were euthanized by CO_2_ inhalation.

### Flow cytometry

Cervical skin draining lymph nodes (sdLN) were removed from the mice and immediately homogenized manually through a 100 μm cell strainer (Fisher Scientific) into a 50 mL tube with the end of a plunger from a 3 mL syringe. The strainer was washed with wash medium (2% vol/vol FBS/PBS) and the strained cells were centrifuged at 400g for 5 minutes at 4°C. Lymph node cell samples were stained with Zombie NIR viability dye (Biolegend; 1:500) to exclude dead cells, followed by Fc-receptor blocking and cell-surface staining with specific antibodies. The cells were analyzed using either LSR Fortessa^TM^ (BD) or Cytek^®^ Aurora (CYTEK) flow cytometers. Data was obtained using either FACSDiva^TM^ (BD) or SpectroFlo® (CYTEK) software and was further analyzed using FlowJo^TM^.

### Lymphocyte stimulations

After tissue harvest, ILC stimulations were performed by incubating 0.5-1×10^6^ cells for 4 hours at 37°C in stimulation media (DMEM with 5% fetal bovine serum, 1% penicillin/streptomycin, 2 mM L-glutamine, 50 ng/mL Phorbol 12-myristate 13-acetate (PMA), 100 ng/mL ionomycin, 5 ug/mL Brefeldin A (BFA), 2 uM monensin). T cell stimulations were performed by first coating a 96-well plate with 5 μg/mL anti-mouse CD3 (Biolegend) in 50 μL/well PBS overnight the day before tissue harvest. The following day, 0.5-1×10^6^ cells were resuspended in 50 μL/well T cell stimulation media (5 μg/mL anti-mouse CD28 (Biolegend), 5 ug/mL BFA), 2 μM monensin) and incubated for 20 minutes hours at 37 C. The cells were then transferred to the anti-mouse CD3 coated plate and incubated for 4 hours at 37 C. After stimulation, cells were washed in wash medium, fixed, and stained for surface and intracellular markers as described for unstimulated cells.

### Splenocyte chimeras

Spleens were harvested from donor WT B6 mice and immediately homogenized manually through a 100 μm cell strainer (Fisher Scientific) into a 50 mL tube with the end of a plunger from a 3 mL syringe. The strainer was washed with wash medium (2% vol/vol FBS/PBS) and the strained cells were centrifuged at 400g for 5 minutes at 4°C followed by treatment with RBC lysis buffer for two minutes and two wash steps using 2 volumes of wash medium. Cells were counted, and 5 million splenocytes were injected intraperitoneally into each recipient mouse. Experiments were performed 4 weeks following splenocyte add-back to allow immune reconstitution.

### Bone marrow chimeras

Recipient mice were provided with antibiotic water, consisting of 5 mL of Sulfatrim (sulfamethoxazole/trimethoprim) added into 200 mL of drinking water, for one week starting from the day prior to irradiation (day −1). On day 0, recipient mice were irradiated with 950 cGy using the X-RAD 320 (Precision X-Ray). BM was harvested from donor mice femurs and tibias and treated with RBC lysis buffer (Sigma-Aldrich) for two minutes. BM cells were transferred into a 15 mL conical tube through a 70 μm cell strainer (Fisher Scientific) and the cell strainer and cells were washed with 2% (vol/vol) FBS/PBS. The concentration of living cells was determined using a Cellometer Auto 2000 (Nexcelom Bioscience) with ViaStain^TM^ AOPI Staining Solution (Nexcelom Bioscience). Recipient mice received the same number of cells, at 1 x 10^7^ live bone marrow cells per mouse, through retroorbital injection within 24-hour after irradiation. Recipients were given 8 weeks for immune reconstitution after BM transplantation before experimental use.

### Cryopreserving sdLN cells for sequencing

*Rag1*^Cre^::*Rosa26*^LSL-tdRFP^ mice were treated with 2 nmol calcipotriol (MC903, Tocris Bioscience) in 10 μL of 100% ethanol (EtOH) vehicle, or vehicle alone, on the bilateral ear skin daily for 7 days to induce AD-like inflammation. The next day, cervical sdLN were harvested and immediately homogenized manually through a 100 μm cell strainer (Fisher Scientific) into a 50 mL tube with the end of a plunger from a 3 mL syringe. The strainer was washed with wash medium (2% vol/vol FBS/PBS) and the strained cells were centrifuged at 400g for 5 minutes at 4°C. Next, cells were incubated with biotinylated antibodies (anti-mouse CD3e, CD19, CD11b; 1:300; Biolegend) in 100 μL of wash buffer for 20 minutes at 4°C, followed by two washes in 2 volumes of wash buffer. Next, no more than 10^7^ cells were incubated with Streptavidin MicroBeads (Miltenyi) in 500 μL separation buffer (0.5% w/v BSA in PBS; BSA and PBS from Sigma) at 4°C for 20 minutes, then added to LD columns (Miltenyi) pre-equilibrated with separation buffer and loaded in a QuadroMACS™ Separator (Miltenyi) for negative cell selection. Remaining cells were eluted in 1 mL separation buffer and cells were centrifuged at 400g for 5 minutes at 4°C, followed by resuspension in freezing buffer (10% DMSO, Invitrogen; 20% FBS in DMEM, Sigma) and slow freezing to −80°C in a CoolCell™ LX (Corning) device.

### Processing cryopreserved cells for multiome

Cryopreserved sdLN cells were processed as recommended by the 10X Genomics DemonstratedProtocol_NucleiIsolation_ATAC_GEX_Sequencing_RevC_(CG000365) instructions for primary cells without any modification to the protocol. Briefly, cells were thawed in a 37°C water bath followed by dilution into media (RPMI + 15% FBS, Sigma) and centrifugation at 400g for 5 minutes at 4°C. For each final sample (EtOH vehicle- or MC903-treated), cells were pooled from sample from 3 individual mice. Cells were resuspended in PBS + 0.04% BSA (Sigma) and passed through a 40 μm Flomi strainer (Bel-art) followed by determination of cell concentration using the using Cellometer Auto 2000 (Nexcelom Bioscience) with ViaStain^TM^ AOPI Staining Solution (Nexcelom Bioscience). Cells were centrifuged 5 minutes at 4°C and supernatant removed. Lysis Buffer (Tris HCl base with 0.1% Tween-20, 0.1% NP-40, 0.01% digitonin, 1 mM DTT, and 1 U/μL Protectors RNase inhibitor, Sigma; full recipe in 10X protocol) was added, cells mixed 10x, and incubated on ice for 3 minutes. Nuclei from lysed cells were centrifuged at 400g for 5 minutes at 4°C and washed in 1 mL Wash Buffer (Lysis Buffer, but without NP-40 or digitonin). The wash step was repeated two more times. Nuclei concentration was determined as for cell concentration using the Cellometer and ViaStain^TM^ solution. The AOPI staining indicated 97-99% lysis efficiency of the cells. We manually confirmed nuclei count using a Bright-Line™ hemacytometer (Hausser Scientific™). Nuclei were centrifuged at 400g for 5 minutes and resuspended in a volume of 1X Nuclei Buffer (10X Genomics) to yield roughly 4,000 nuclei/μL. We then immediately proceeded to the 10X Chromium Next GEM Single Cell Multiome ATAC + Gene Expression pipeline.

### Multiome library construction and sequencing

Multiome 3v3.1 GEX and ATAC libraries were prepared as recommended by 10X Genomics protocol Chromium_NextGEM_Multiome_ATAC_GEX_User_Guide_RevD ((CG000338). For sample preparation on the 10x Genomics platform, the Chromium Next GEM Single Cell Multiome ATAC + Gene Expression Reagent Bundle, 16 rxns PN-1000283, Chromium Next GEM Chip J Single Cell Kit, 48 rxns PN-1000234, Single Index Kit N Set A, 96 rxns PN-1000212 (ATAC), Dual Index Kit TT Set A, 96 rxns PN-1000215 (3v3.1 GEX), were used. The concentration of each library was accurately determined through qPCR utilizing the KAPA library Quantification Kit according to the manufacturer’s protocol (KAPA Biosystems/Roche) to produce cluster counts appropriate for the Illumina NovaSeq6000 instrument. GEX libraries were pooled and run over 0.05 of a NovaSeq6000 S4 flow cell using the XP workflow and running a 28×10×10×150 sequencing recipe in accordance with manufacturer’s protocol. Target coverage was 500M reads per sample. ATAC libraries were pooled and run over 0.167 of a NovaSeq6000 S1 flow cell using the XP workflow and running a 51×8×16×51 sequencing recipe in accordance with manufacturer’s protocol. Target coverage was 250M reads per sample.

### Multiomic data analysis

The cellranger-arc-2.0.0 (10X Genomics) pipeline was used to generate FASTQ files, gene expression matrices, and ATAC fragment tables for each sample, followed by aggregation using the aggr function. Default settings were utilized, with the exception that we incorporated a custom reference with the sequence for tdRFP (see **Supplemental file S1**) added to the default mouse reference sequence provided by cellranger (refdata-cellranger-arc-mm10-2020-A-2.0.0). Correction for ambient RNA was performed using SoupX^148^ with clustering information provided by the default cellranger outputs. Doublets were removed using Scrublet^149^ with default settings.

Corrected data was then processed using Signac^37^ and Seurat^34–36^. ATAC-seq peaks were identified using MACS2^150^ through the CallPeaks function in Signac. Per-cell quality control metrics were computed using the TSSEnrichment and NucleosomeSignal functions, and cells retained with a nucleosome signal score < 1.5, TSS enrichment score > 1, total RNA counts < 15,000 and > 1,000, total ATAC counts < 75,000 and > 100, percent mitochondrial reads < 5%, and percent ribosomal genes detected <10%. After these filtering steps, 10,304 cells remained. Cells were further filtered by their expression of lineage defining markers similar to the negative selection step during sample processing. Cells with detectable transcripts for *Cd3d*, *Cd3e*, *Cd3g*, *Cd4*, *Cd19*, *Cd8a*, and *Itgam* were removed. This left 2,034 remaining cells for further analysis.

The SCTransform function of Seurat was used to normalize RNA counts. We performed integration of the two samples using the RNA assay to correct for batch effects and treatment in the initial clustering using the default parameters for the functions SelectIntegrationFeatures, FindIntegrationAnchors, and IntegrateData. The integrated data was used for PCA (25 dimensions) and UMAP reduction for the RNA assay alone. With default parameters in Signac, we used TFIDF to normalize ATAC peaks and latent semantic indexing (LSI) to reduce the dimensionality of the ATAC data. We constructed a UMAP of the ATAC data alone using the LSI reduction (dimensions 2-25). To construct a joint graph and UMAP using equal weighting from the RNA and ATAC assays, we used the FindMultiModalNeighbors function of Seurat/Signac using default parameters (RNA dimensions 1-25, ATAC dimensions 2-25). We used a resolution of 0.1 to identify clusters with the FindClusters function in Seurat/Signac. Cell types were assigned based on manual curation of marker genes. Initially, 7 clusters were identified, but two highly similar lymphocyte clusters were merged for a total of 6 cell types.

The inferred Gene Activity (GA) assay from the ATAC-seq data was calculated using default parameters of the GeneActivity function in Signac. FindAllMarkers was used to identify top markers by cluster for both RNA gene expression data (GEX) and GA, with setting adjustments including min.pct = 0.20 and logfc.threshold = 0.25. The differentially accessible (DA) open chromatin assay was calculated in Signac with the FindMarkers function on the ATAC-seq peaks assay (called using MACS2 as above). The differential test used was ‘LR’ (logistical regression, as suggested for snRNA-seq^151^). The total number of ATAC fragments was used as a latent variable to mitigate effect of differential sequencing depth. Given the sparsity of the data, the min.pct parameter was set to 0.02. After identifying the top differentially accessible peaks for each cluster, the gene closest to each peak was determined using the ClosestFeature function in Signac. Results were filtered for genes within 10^5^ base pairs of the corresponding peak. The filtered gene lists were used for the “DA” assay as markers of each cluster (top 25) and an expanded list for the ILC2 cluster (top 100). Venn diagrams were calculated using BioVenn/BioVennR^152^.

Gene set enrichment analysis was performed and visualized using ClusterProfiler^153^. For GSEA on steady state ILC2 DEGs between fate mapped states, we opted to use more permissive filtering parameters instead of default parameters. We created the ranked list of DEGs using the FindMarkers function in Seurat with min.pct = 0.1 and logfc.threshold = 0.1. The DEG list from the GEX assay was used to generate the GSEA results. The DEG list from the GA assay did not yield any significant GSEA results. The ClusterProfiler function gseGO was used to analyze the ranked DEG list using the paramters minGSSize = 50, maxGSSize = 500, pvalueCutoff = 0.05.

A motif matrix was constructed from the ATAC data Granges using the “CORE” collection and “vertebrates” taxonomy group from the JASPAR2022 position weight matrix set and the mm10 reference genome. Per cell transcription factor motif activity was calculated with chromVar^94^ using the motif matrix and MACS2 called peaks. Transcription factor motifs were identified in differentially accessible chromatin using the FindMotifs function in Signac.

The correlation coefficients, or gene to peak links (GPLs), between gene expression and accessibility of each peak were calculated for all peaks within 10^6^ base pairs of the transcription start sites for all detected genes using the LinkPeaks function of Signac with min.cells = 2. GPLs were filtered by gene for the curated ILC2 and Th2 gene sets. Since multiple genes can be linked to one peak by GPL analysis, finding intersections of GPLs in set analysis would result in counting some epigenomic regions multiple times. Thus, for set analysis, we eliminated GPLs with redundant peaks. Then, we used each list of non-redundant peaks as input sets to generate UpSet plots and lists of intersecting peaks between states (Rag1 fate map positive or negative; AD-like disease or steady state) using UpSetR^154^. Coverage plots of the single cell multiomic data, including open chromatin, peaks, and links (GPLs), were plotted using the CoveragePlot function in Signac.

### Data and code availability

Sequencing data have been deposited at GEO and accession numbers are listed in the key resources table. Aggregated data are supplied in the supplemental file. All data reported in this paper will be shared by the lead contact upon request. This paper does not report original code. Any additional information required to reanalyze the data reported in this paper is available from the lead contact upon request.

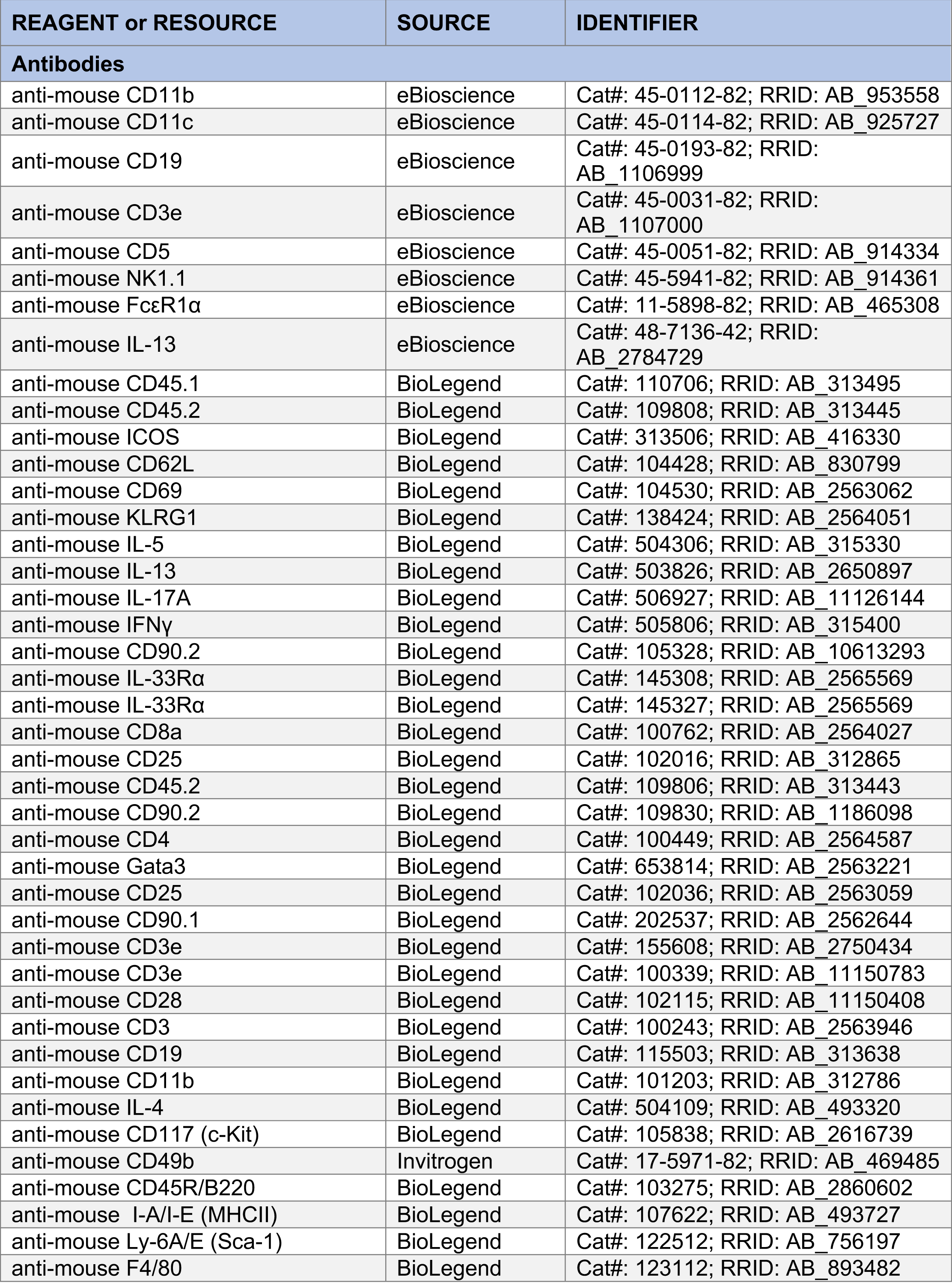
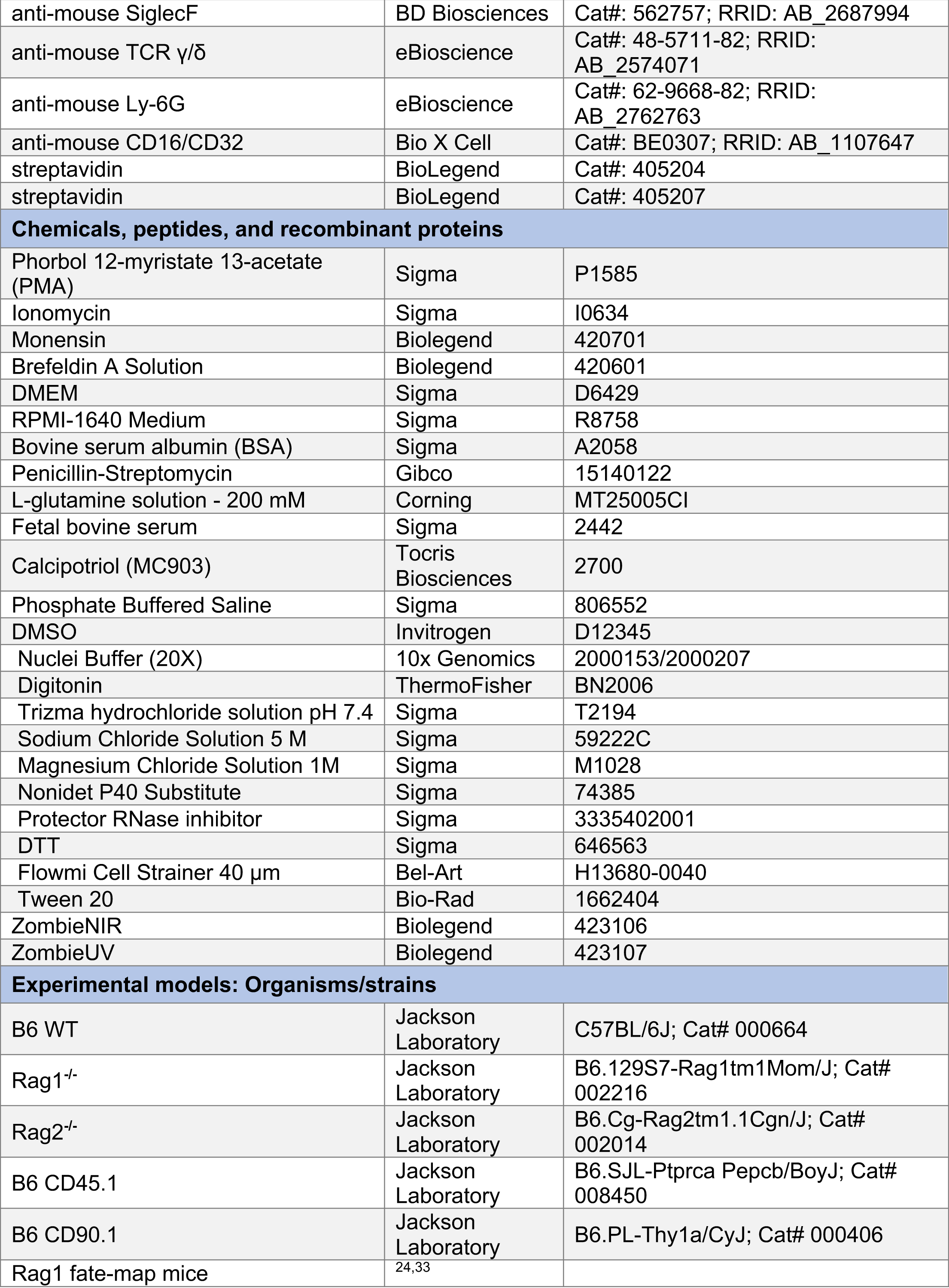
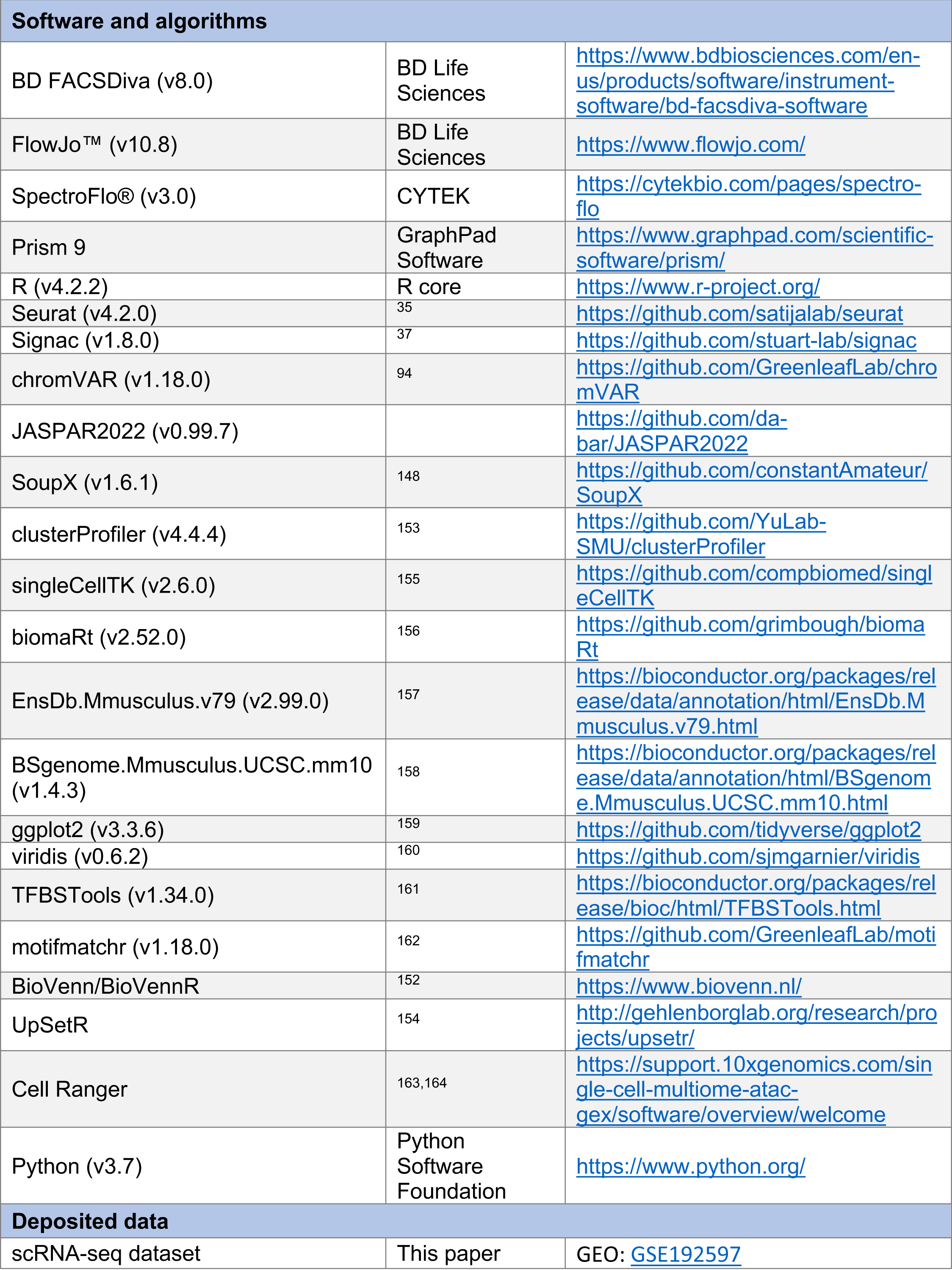
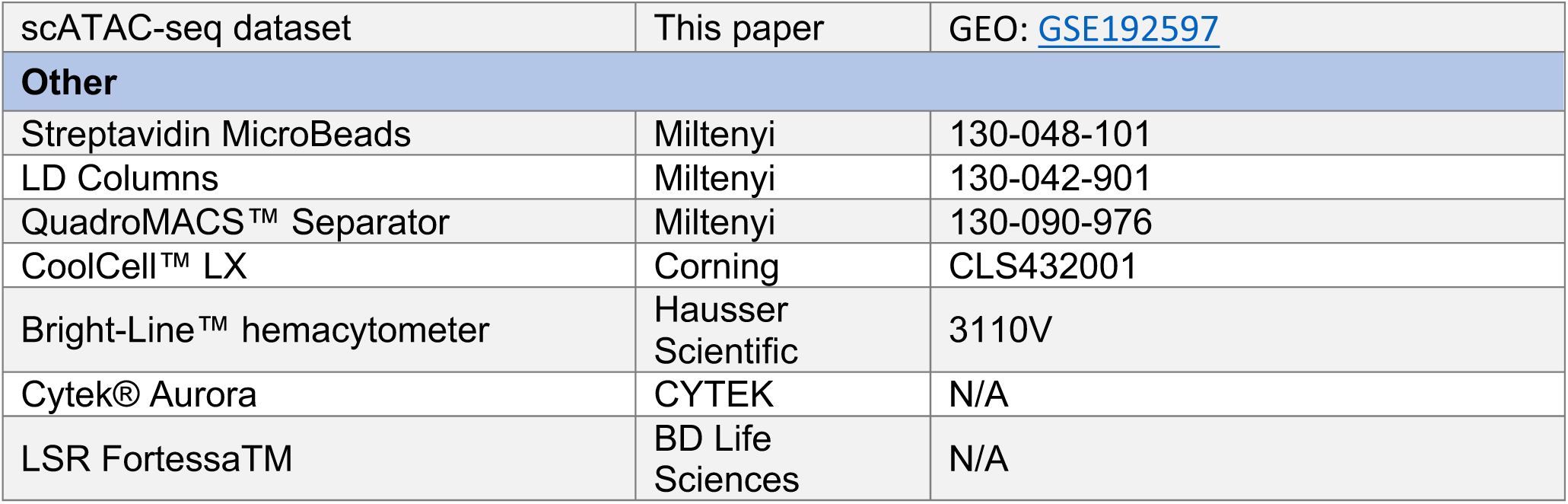

